# SPrUCE: Utilizing Ultraconserved Elements of DNA for Population-Level Genetic Diversity Estimation

**DOI:** 10.1101/2025.11.14.688492

**Authors:** Daira Melendez, Ali Osman Berk Şapci, Vineet Bafna, Siavash Mirarab

**Affiliations:** Bioinformatics and Systems Biology Graduate Program, UC San Diego, San Diego, CA, USA; Department of Computer Science and Engineering, UC San Diego, San Diego, CA, USA; Department of Electrical and Computer Engineering, UC San Diego, San Diego, CA, USA

**Keywords:** ultraconserved elements, population genomics, nucleotide diversity, conservation genetics

## Abstract

Ultraconserved elements (UCEs) provide ideal candidates for targeted sequencing and cost-effective acquisition of genome-wide data. While UCEs have been widely used in phylogenetic studies to reconstruct evolutionary relationships, their use in population-level research has been limited. This limited application stems from uncertainty over whether UCEs can capture the levels of genetic variation needed to answer population genomic questions central to ecology and biodiversity research. The concern is that, by definition, UCEs are highly conserved and may therefore lack sufficient within-species variation. The more variable flanking regions (400–750 bp from the UCE core) contain informative polymorphisms, though diversity decreases near the core. Thus, any naive estimator of genetic diversity that ignores this conservation will have an underestimation bias. In this paper, we introduce *SPrUCE: Sigmoid Pi requiring UCEs*, a reference-free method that estimates nucleotide diversity *π* from aligned UCE data. SPrUCE corrects underestimation bias by modeling the change in diversity away from the UCE core using a Gompertz function. The model accounts for the bias introduced by the conserved core and allows for more accurate per-site diversity estimates. We tested SPrUCE on UCE alignments from a range of taxa, including invertebrates and vertebrates (finches, honeybees, sheep, and smelt). SPrUCE produces diversity values consistent with whole-genome derived estimates that require an assembled reference. It is fast, scalable, and effective even with missing data. Its modeling approach enables accurate population-level assessments of genetic diversity, offering a new and reliable option for conservation and population genetics.

## 1 Introduction

Measuring genetic diversity is a prerequisite for many biological analyses, including the study of evolutionary processes such as population differentiation and speciation, as well as biodiversity monitoring and ecological assessments (Hughes et al., 2008). The recent and rapid loss of biodiversity (IPBES, 2019) has created an urgent need to monitor “essential biodiversity variables,” several of which can be quantified by measuring genetic diversity within populations (Pereira et al., 2013). Rapid estimation of genetic variation is now recognized as a critical tool for successful conservation (Kardos et al., 2021), particularly when diversity is measured genome-wide (Supple and Shapiro, 2018). At the same time, the increasing scale of biodiversity decline across taxa demands methods that enable genetic monitoring at shallow population-level scales necessary for conservation and management of threatened populations.

Genome-wide diversity is often defined with *θ* = 4*µN*_*e*_, a measure of the effective size of the population scaled by the mutation rate (Watterson, 1975). It is often estimated using methods such as nucleotide diversity (*π*) and Watterson’s theta estimate, which are computed from a matrix of nucleotide polymorphisms (a SNP matrix) in a sample of the population. The SNP-matrix is generated by mapping sequenced fragments (typically at high coverage 30*×*) to an assembled reference genome. Despite the drop in the cost of whole genome sequencing (WGS), this method remains costly for biodiversity studies, which need rapid and repeated sampling, and an assembled reference. A more affordable option is to sequence samples at low coverage and use genotype likelihood modeling (e.g., ANGSD Korneliussen et al., 2014) to distinguish between sequencing errors and real substitutions. This option also has drawbacks. It requires an assembled reference, which is not available for much of the Earth’s biodiversity. Second, for very low sequence coverage (e.g., *<* 4*×*), its estimates become inexact and sensitive to stringency parameters. Especially in conservation genetics, where researchers often work with degraded or limited DNA (Bi et al., 2013), it may only be possible to obtain uneven and low coverage. Moreover, for large genomes, even low coverage sequencing can be expensive. Finally, the computational expense cannot be ignored, as it often requires hours to days to process data using tools like GATK (McKenna et al., 2010) or ANGSD, limiting their feasibility in field-based research.

Recognizing that less data may be sufficient for estimating diversity, many researchers instead sequence predefined loci using methods such as RAD-seq (Davey et al., 2011), mitochondrial capture (Liu et al., 2016), or targeted capture (Jones and Good, 2016). By targeting a small fraction of the genome, these methods improve cost-effectiveness, and they can also reduce the sequencing of off-target DNA from unintended sources, which can be a major issue (Jones and Good, 2016). However, sequencing a non-random subset of loci often leads to biases such as underestimation of diversity (Arnold et al., 2013) and estimates that do not represent the entire genome (Galtier et al., 2009). For example, allele dropout in RAD-seq data has been shown to systematically underestimate diversity (Arnold et al., 2013), necessitating computational modeling to correct these biases.

An increasingly adopted source of data is targeted capture of ultraconserved elements (UCEs). UCEs are small DNA segments that exhibit exceptionally high sequence conservation (e.g., *>*97% identity; Bejerano et al., 2004; Cummins et al., 2024) within divergent lineages, including placental mammals, birds, reptiles, insects, and fish. This extreme conservation enables their utility as universal markers across broad phylogenetic scales (Faircloth et al., 2013). The sequence conservation is helpful because it simplifies designing baits that hybridize with a wide range of species, eliminating the need to design species-specific baits. The conserved cores serve as anchors to also capture the flanking regions (Faircloth et al., 2012), which show increased genetic variability (Fig. 1a,b). In addition to ease of sequencing UCEs are distributed across the genome (Cummins et al., 2024) and can therefore be representative of the entire genome. UCEs were initially proposed for phylogenetic reconstruction and they are now adopted across a wide range of taxonomic groups (e.g., Faircloth et al., 2015; Miles Zhang et al., 2019; Esselstyn et al., 2017; Alfaro et al., 2018; Starrett et al., 2017).

**Figure 1.**
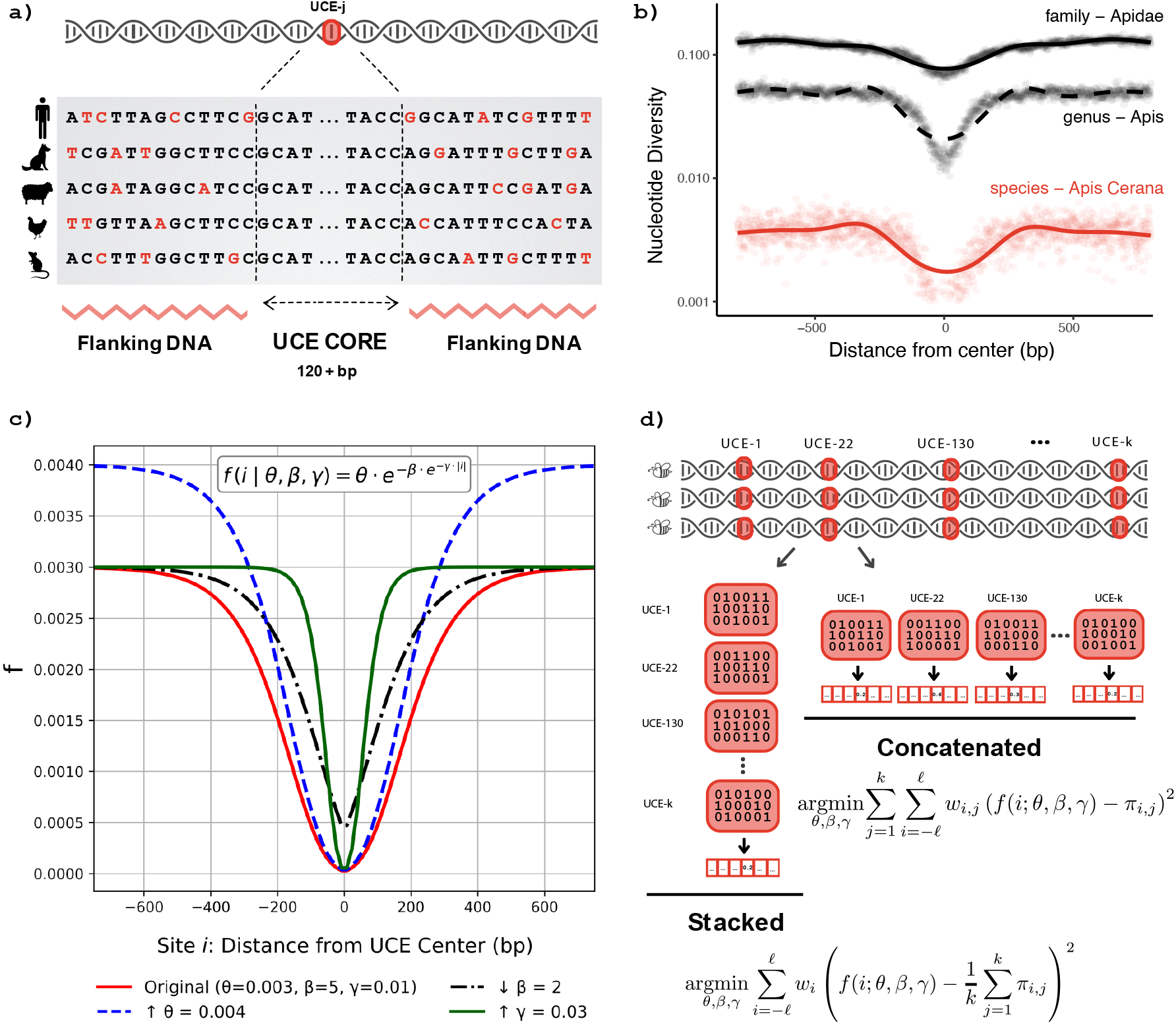
The logic of SPrUCE demonstrated by (a) cartoon of Ultraconserved element (UCE) across species, with variable flanking region, (b) UCE heterozygosity across honeybee taxonomic ranks, computed using Eq. (1), (c) Gompertz function with four sets of parameters, and (d) the two modes of the SPrUCE method: Stacked (aggregating frequencies across UCEs) and Concatenated (analyzing individual UCEs jointly).

Beyond phylogenetics, several authors have used UCEs for comparing recently differentiated subspecies or even measuring population genetic measures of diversity (Stiller et al., 2021; Winker et al., 2018; Smith et al., 2013; Manthey et al., 2016; Oswald et al., 2016). What they all show is that while within-species genetic diversity is low and further reduced within the core region, as one moves to the flank, there are enough substitutions to make meaningful statements about population dynamics (see Fig. 1b for an example among bees), especially population structure. While these studies show a signal, especially to differentiate subpopulations, less attention is paid to how to account for the bias they introduce.

A key difficulty in using UCEs for measuring population-level genetic diversity is that, by definition, they are highly conserved, which can therefore bias the estimates. Simply using the aligned UCE sites as if they represent the genome will lead to underestimation. A simple alternative would be to use flanks only; however, due to recombination dynamics (and perhaps other factors related to the biology of UCEs), as we transition from the (highly conserved) core to the (presumably neutral) flanks, the change in genetic diversity is gradual (Fig. 1b), making it non-trivial to know where to cut. A more theoretically justifiable method is to model the diversity as a function of distance from the core and use the model to compensate for reduced variation within UCEs.

The goal of this paper is to present a modeling framework that allows unbiased estimates of population-level genetic diversity. We focus specifically on estimating nucleotide diversity (*π*) (Nei and Li, 1979), which measures the average pairwise nucleotide differences per site across sequences sampled from a population. For a population evolving according to the neutral Wright Fisher model, E[*π*] = *θ* (Tajima, 1983). To model the observed patterns of diversity (Fig. 1b), we explore sigmoid-based functions, ultimately selecting the Gompertz function (Fig. 1c) due to its biological interpretability, the relative ease of fitting its three parameters, and its empirical fit to our data. We implement this approach in a tool called SPrUCE (Sigmoid Pi requiring UCEs) with a streamlined processing pipeline that follows the standard Phyluce protocol (Faircloth, 2016) and outputs estimates of nucleotide diversity *π*. SPrUCE is designed for scalability and can complete analyses in a few seconds to minutes. We examine the accuracy and versatility of SPrUCE in a series of analyses on UCEs extracted from real genomic data (where we can compare against WGS estimates) and real targeted capture UCE data. Results show that SPrUCE dramatically reduces bias and matches WGS estimates, enables estimations from relatively small flanking regions, and scales easily.

## 2 Materials and Methods

### 2.1 SPrUCE Algorithm

The input to SPrUCE is a set of *k* UCE alignment files, one for each UCE locus. Each file describes a multiple sequence alignment with rows corresponding to individuals and columns corresponding to sites centered around the UCE core (Figure 1d). In the following, we will use ‘UCE’ to refer to individual UCE alignments, and ‘sites’ refer to specific nucleotide positions within those alignments (Figure 1a). Popular pipelines such as Phyluce (Faircloth, 2016) directly produce the required input. SPrUCE’s output is an *estimate* of nucleotide diversity, *π*.

#### 2.1.1 Correcting biased estimates of diversity using the Gompertz model

Let *s*_*i*_ be the count of individuals carrying the minor allele at site *i*, and *n*_*i*_ be the number of individuals. Then,

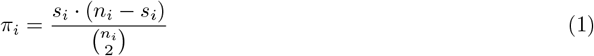

gives a per-site estimate of nucleotide diversity; these per-site estimates can be averaged across *L* sites to get a genome-wide estimate: 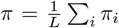. This estimator can be applied to any subset of sites, provided that those sites can be assumed to follow the same distribution as the rest of the genome. Under neutral evolution (i.e., no selection) and a large constant population size, *π* is a valid estimator of *θ* (Tajima, 1983). Note that there is a Phyluce program that outputs a “substitution frequency” metric, which, as we show in the Supplementary material, is approximately *π*_*i*_*/*2 even if we ignore the UCE conservation.

Applying the standard *π* estimator directly to ultraconserved elements (UCEs) can result in a underestimation of the diversity parameter *θ* due to the (presumed) presence of selection. The strong conservation of the sequences near the UCE core (Fig. 1a) is a signature of negative selection and reduced *π* (Katzman et al., 2007; Cummins et al., 2024). We can represent UCE alignments as analogous to a SNP matrix, where rows correspond to individuals and columns to positions of nucleotides (Fig. 1d). As we move from the center to the flanking regions, the diversity increases, eventually approaching a plateau (Fig. 1b). These plateaus presumably correspond to genome-wide levels of diversity with little or no linkage to the conserved core. Clearly, just averaging Equation (1) across all sites of a UCE will lead to a biased estimator that underestimates *π*. To reduce such a bias, one has to correct for the conservation at the core.

Our proposed approach is to model the change in *π*_*i*_ across a UCE locus using the Gompertz function (Gompertz, 1825) and to estimate *θ* by fitting parameters of Gompertz to the data. Gompertz is a sigmoid-shaped function and can reach a parametrized asymptote with adjustable rates and intercepts (Fig. 1c). The asymptote parameter effectively models the plateau level in the distant flanking region, which we assume represents *θ*. More precisely, we assume: 𝔼[*π*_*i*_] = *f* (*i*; *θ, β, γ*) where

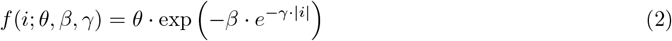

Here, the variable *i* corresponds to the positional distance from the UCE core, measured in base pairs, and can be positive or negative (the absolute value ensures symmetry around position 0, which is the UCE center).

#### 2.1.2 Interpreting parameters of the Gompertz function

The Gompertz model was selected empirically for its flexibility in capturing the characteristic pattern of UCEs, where diversity is minimized in the highly conserved core bases and increases symmetrically toward the flanking regions. Compared to the generalized logistic function (which has six parameters), Gompertz has the advantage of having fewer parameters and being easier to fit. One can attempt to use even simpler models, such as a piecewise linear model, but such models do not fit the data as well, as we will see in the results. Moreover, Gompertz’s three parameters have biological interpretations.

We can provide an interpretation of the parameters *β, γ* by assuming that the UCE region is under negative selection at the center of the UCE with the following properties: selection coefficient *s*, and per nucleotide rates of deleterious mutation, *u*, where *u ≪ s*, and recombination *r* (*r ≪ s*). Then, following Hudson and Kaplan (1995) (cf. Eqn. 3), the local diversity at distance *x* from the center is given by

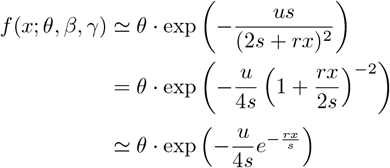

provided *x* is small enough that 2*rx ≪ s*. Comparing with the Gompertz equation, parameters 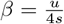 and 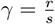 are governed by the strength of selection and the rates of deleterious mutations and recombination in the region. While the original HK95 equation could also be used as a model, we show in this paper that the approximate Gompertz function provides a better empirical fit across a wider range of flank sizes.

#### 2.1.3 Fitting parameters of Gompertz

To fit the Gompertz function parameters *θ, β, γ*, we suggest two estimators: Stacked, which combines all positions with the same index across all UCEs into one data point, and Concatenated, which treats each site of each UCE as an independent estimator (Fig. 1d).

The Stacked mode assumes patterns of change in diversity across loci are similar and thus combines all UCE loci into a single matrix for calculating the diversity metric. Let 2*ℓ* + 1 be the number of sites (*ℓ* on each side of the center, indexed at 0) that each of our *k* UCE alignments is assumed to have. We use a weighted least squares estimator:

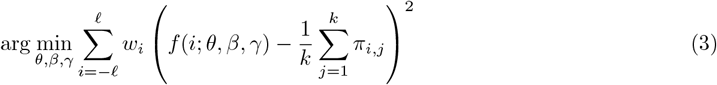

where *π*_*i,j*_ is the diversity of site *i* from locus *j*, defined as in Equation (1), with a caveat described below; *f* (*i*) is the Gompertz function given in Equation (2). The weights *w*_*i*_ are used to adjust the impact of different positions on the estimator, as explained below.

In the Concatenated mode, Gompertz parameters are fit to all sites across all UCE loci and allows a more fine-grained weighting scheme. The optimization goal is to find:

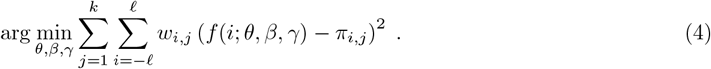

where weights *w*_*i,j*_ can now change across loci. Note that while the Stacked optimization problem is over *O*(*ℓ*) terms, the Concatenated problem has *O*(*ℓk*) terms, making it more time-consuming to solve.

Both modes need ways to deal with length heterogeneity across UCEs as well as missing individuals, insertion and deletions, and alignment issues. When a locus *i* has fewer than 2*ℓ* + 1 sites, the sum in Equation (4) is trivially adjusted to only include the available sites. Similarly, in the Stacked mode, the average 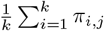 is adjusted to only include available loci for each position *i*.

Beyond length heterogeneity, we also need to account for missing individuals and gaps. Intuitively, sites with higher coverage should lead to lower variance in *π*_*i,j*_, and are hence more reliable and should contribute more strongly to the final estimate. By Aitken’s theorem, the best linear unbiased estimator (BLUE) can be achieved under certain conditions by weighting each term in the least squares estimator by the inverse of the variance of the terms (Aitken, 1936). For *n* individuals, Tajima (1983) has calculated the variance of the *π* estimate to be

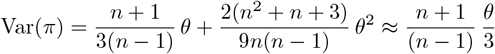

where the approximation holds for sufficiently small *θ*. Thus, to obtain Aitken’s BLUE estimator, we can simply set:

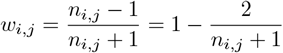

where *n*_*i,j*_ (locus/site coverage) is the number of individuals for which locus *j* has a non-gap character in its alignment at site *i*. When many individuals are missing, the small *n*_*i,j*_ ensures the site gets a low weight (Fig. S1). Thus, sites with more missing data or indels are down-weighted, and more reliable positions are given more weight.

As we move towards the flanking regions, we get to indices where very few loci have any information (Fig. S2). These sites are often alignment artifacts and unreliable. In the Stacked mode, we define weights using *n*_*i*_, which is the sum of *n*_*i,j*_ over all loci *j*. However, since *n*_*i*_ is a large value, all weights are close to 1 and therefore less impactful (Fig. S1). To ensure that only reliable data contribute to the diversity metric, positions with base pair coverage (*n*_*i*_) below a certain threshold (*δ*) are excluded in the Stacked mode. To identify *δ*, instead of using a fixed value, we use a data-driven approach. We empirically observed that the distributions of base pair coverage across sites are left-skewed with a long tail of sites with low coverage (Fig. S2). Thus, we seek to identify the inflection point of the coverage distribution as *δ*. However, since coverage values are discrete and the true probability mass function is unavailable, we use the empirical cumulative distribution function (ECDF) as a proxy. In particular, to reduce noise, we fit a spline function to the ECDF after applying log-transformation (Fig. S2). Finally, we set *δ* to the coverage value near where the second derivative changes its sign.

#### 2.1.4 Implementation details

We solve both Stacked and Concatenated least-squares optimization problems using a bounded nonlinear optimizer implemented with the `trust-constr` method from the `scipy` library (Virtanen et al., 2020). To ensure biologically realistic fits, we set the initial parameters for the Gompertz function to [0.001, 10, 0.009], corresponding to *θ, β*, and *γ*, respectively. Upper and Lower bounds are defined as *θ ∈* [0, 0.05], *β ∈* [0.1, 30], and *γ ∈* [0.004, 0.03]. These ranges were chosen based on observed diversity levels in empirical datasets and typical Gompertz curve behavior when modeling nucleotide diversity near UCEs. For example, *θ* values are generally small (often *<* 0.01) in species populations, while the bounds on *β* and *γ* ensure the curve transitions smoothly within the flanking region and reaches a plateau consistent with biological expectations.

We implemented SPrUCE in Python by extending an existing Phyluce program (Faircloth, 2016). SPrUCE combines the new optimization with several data processing steps (see Algorithm S1). Since we focus on population genetic scales, we only count biallelic sites, removing the multi-allelic sites.

### 2.2 Evaluations

#### 2.2.1 Datasets

We tested SPrUCE on two types of datasets: We first evaluated the accuracy of SPrUCE on four genomewide datasets with multiple species and populations where we could obtain a reference estimate based on genome-wide data, and compare it with SPrUCE run on UCEs curated from the same data. We then evaluated SPrUCE on two UCE datasets for which no ground-truth estimates are available.

##### Whole Genome Datasets

Raw sequencing reads were obtained from NCBI and previously published whole-genome sequencing (WGS) datasets, including honeybees (*Apis cerana*), finches (*Certhidea fusca, Geospiza conirostris*, and *Pinaroloxias inornata*), sheep (*Ovis aries*), and smelt fish (*Sillago sinica*). See Tables S1 to S4 for read IDs and details. For the honeybees (*Apis cerana*), we used 20 individuals from two sub-populations, Niupeng and Zhongshui, from the Chinquai region of China as studied by Wang et al. (2024). We utilized the Hymenoptera v2 UCE bait set containing 2,590 UCEs (Branstetter et al., 2017). For the smelt (*Sillago sinica*) dataset by Zhao et al. (2023), we used 43 individuals from three subpopulations: Dongying, Qingdao, and Wenzhou. We used the Acanthomorph UCE bait set containing 1,314 UCEs (Alfaro et al., 2018). For the finch dataset of Lamichhaney et al. (2015), we used 46 individuals from three species: *Certhidea fusca, Geospiza conirostris*, and *Pinaroloxias inornata*, sampled from different island populations (Cristobal, Espanola, Genovesa, and Cocos Island). We used the Tetrapods probe set, targeting 5,060 UCEs (Faircloth et al., 2012). We also applied the Tetrapods bait set for the sheep (*Ovis aries*) dataset of Shi et al. 2023 on 20 individuals, 10 from each of two populations: heritage Oula sheep and domesticated Panou sheep. In total, our WGS UCE dataset includes 129 individuals from 12 populations, across six species, with three different UCE bait sets.

Whole-genome sequencing (WGS) reads were processed prior to UCE extraction (Fig. S3). Raw reads were quality filtered (QC *>* 20) and adapter-trimmed using BBMap (Bushnell, 2014). De novo assemblies were generated for each individual using MEGAHIT v1.2.9 (Li et al., 2015) and assembly quality was assessed using BBMap stats. Assembled genomes were used as input and processed through the standard Phyluce pipeline (v1.7.1) (Faircloth, 2016) to identify and extract UCE loci and their flanking regions. For each UCE, we extracted 400 bp and 750 bp of flanking sequence on both sides of the conserved core. These fragment sizes were selected to evaluate the effect of flanking region length on nucleotide diversity estimates: 400 bp reflects length typical of UCE capture, while 750 bp approximates longer UCE fragment recovery (McCormack et al., 2016). UCE loci were aligned across individuals using MAFFT (Katoh and Standley, 2013), generating one alignment per UCE locus for both flanking-length datasets. The resulting UCE alignments were used as input for SPrUCE analyses. Tables S1 to S4 lists the number of UCE loci recovered per individual, and individual sample identifiers.

##### UCE Datasets and Effective Population Size (*N*_*e*_) Estimation

We used two UCE MAFFT alignment datasets from previously published studies. Oswald et al. (2016) applied UCE data for Willets (*Tringa semipalmata*) and estimated nucleotide diversity and effective population sizes for the two subspecies *T. s. semipalmata* and *T. s. inornata*. Similarly, Winker et al. (2018) used UCE loci to infer population parameters, including nucleotide diversity, effective population size (*N*_*e*_), divergence times, and gene flow between snow buntings (*Plectrophenax nivalis*) and McKay’s buntings (*P. hyperboreus*). We used the UCE alignments provided by these studies and applied SPrUCE to estimate nucleotide diversity (*π*) directly from the alignments.

In addition to estimating *π*, we calculated effective population size (*N*_*e*_) to provide demographic context and evaluate whether our genetic diversity estimates align with previously published values. *N*_*e*_ was calculated using the standard *θ* = 4*N*_*e*_*µ*, where *µ* is the mutation rate per site per generation. For passerines (*Plectrophenax* spp.), we used a mutation rate of 6.75 *×* 10^*−*10^ substitutions/site/year and a generation time of 2.7 years following Winker et al. (2018). For shorebirds (*Tringa semipalmata*), we used a mutation rate of 2.59 *×* 10^*−*10^ substitutions/site/year based on Oswald et al. (2016) and a generation time of 5.9 years based on BirdLife International (International, 2025). Where relevant, we compared *N*_*e*_ values to approximate census population sizes reported by Partners in Flight Partners in Flight (2025) to explore whether genetic diversity reflects contemporary population trends.

Across all datasets, we included a UCE locus if at least three individuals were represented in the alignment.

#### 2.2.2 Validation Criteria

To validate our estimates of nucleotide diversity, we used ANGSD (Analysis of Next Generation Sequencing Data) (Korneliussen et al., 2014) to compare our results to genome-wide estimates. ANGSD maps wholegenome sequencing (WGS) reads to a reference genome to compute genotype likelihoods, infer the site frequency spectrum (SFS), and estimate population genetic statistics such as pairwise nucleotide diversity *π*. We applied ANGSD to adapter and quality-trimmed reads from individuals in each population, mapping them to a representative reference genome to estimate diploid nucleotide diversity genome-wide. These ANGSD-derived *π* values served as an independent benchmark to compare against our diversity estimates derived from UCEs. We compared the Stacked and Concatenated Gompertz-based estimates to genome-wide ANGSD values across flank lengths of 400 bp and 750 bp.

To evaluate the robustness of the Gompertz model in estimating nucleotide diversity from UCEs, we compared its performance against two alternative functions: the Generalized Logistic Function (GLF), with six parameters, and the simpler Piecewise Linear Function (PLF). GLF is a generalization of Gompertz that has more parameters and allows more flexibility in its functional form. PLF is a simpler model with no curve, just two flat lines (one for the asymptote and one for the core) and a line connecting the two. We also compared against the baseline method of using the uncorrected average of diversity across all positions (i.e., simply the mean of Eq. (1) across all sites). We compared how each function performs as the flank size decreases and compared the resulting estimates to genome-wide diversity values derived from ANGSD.

Beyond assessing agreement with ANGSD, we also evaluated the robustness of SPrUCE with respect to (1) the number of UCE loci analyzed, (2) the number of individuals included, (3) dataset completeness (i.e., the proportion of individuals represented per locus), and (4) computational runtime between the Stacked and Concatenated modes. For each dataset, we chose one representative population. To examine the effect of the number of loci on diversity estimates, we generated random subsets of loci in groups of 10, 25, 50, 100, 200, 400, and 800 and performed 10 replicates for each subset size. To evaluate the impact of dataset completeness, we constructed datasets with varying completeness thresholds based on the number of individuals present per locus: 50% completeness (at least half the individuals represented), 80% completeness, and 100% completeness (all individuals present for every locus). This allowed us to explore how increasing completeness (which comes at the cost of fewer loci) impacts SPrUCE estimates. To assess the impact of sample size (i.e, number of individuals), we created subsets that contained randomized 8, 5, and 3 individuals with 10 replicates per subset and compared them to the complete dataset (all individuals present across loci; see Table S5). Finally, we compared the computational runtime of the Stacked and Concatenated and modes in SPrUCE using a single core, quantifying how dataset size and method influence overall analysis time.

## 3 Results

### 3.1 SPrUCE accurately estimates nucleotide diversity

Estimates obtained from UCEs using SPrUCE are highly concordant with ANGSD estimates of *π* from WGS, regardless of the SPrUCE variant used (Fig. 2 and Table 1). SPrUCE estimates exhibit strong Pearson correlation with ANGSD values, with *R*^2^ values ranging from 0.96 to 0.97, with similar correlations obtained for 400bp and 750bp flanks, as well as with Concatenated and Stacked. For every species, the ranking of populations by diversity levels is consistent between ANGSD and SPrUCE, even when the difference between populations is small; for example, honeybees from Zhongshui are slightly more diverse than honeybees from Niupeng for *A. Cerana* in ANGSD (by 3%), a pattern recaptured by every flavor of SPrUCE. In all cases, Kendall’s *τ* and Spearman’s correlations between ANGSD and SPrUCE were 1.0, confirming that both methods rank the populations consistently.

**Table 1:** SPrUCE nucleotide diversity (*π*) estimates from Stacked and Concatenated methods at 400 bp and 750 bp flanks, compared to ANGSD genome-wide estimates.

| Species Group | Population | Size<br>(*n*) | UCE loci<br>400 / 750 | ANGSD<br>$\pi$ | STACKED<br>400 / 750 | CONCATENATED<br>400 / 750 |
| --- | --- | --- | --- | --- | --- | --- |
| *Apis cerana* | Niupeng | 10 | 2414 / 2409 | 0.00544 | 0.00493 / 0.00426 | 0.00481 / 0.00424 |
|  | Zhongshui | 10 | 2410 / 2402 | 0.00560 | 0.00504 / 0.00443 | 0.00488 / 0.00440 |
| *Sillago sinica* | Dongying | 15 | 1005 / 971 | 0.00634 | 0.00621 / 0.00589 | 0.00611 / 0.00580 |
|  | Qingdao | 13 | 1005 / 970 | 0.00639 | 0.00674 / 0.00602 | 0.00653 / 0.00594 |
|  | Wenzhou | 15 | 1005 / 970 | 0.00681 | 0.00711 / 0.00628 | 0.00689 / 0.00617 |
| *Certhidea fusca* | Cristobal | 9 | 4737 / 4729 | 0.00190 | 0.00152 / 0.00146 | 0.00154 / 0.00151 |
|  | Española | 10 | 4751 / 4744 | 0.00063 | 0.00056 / 0.00054 | 0.00052 / 0.00053 |
| *Geospiza conirostris* | Española | 10 | 4739 / 4728 | 0.00237 | 0.00195 / 0.00184 | 0.00193 / 0.00182 |
|  | Genovesa | 9 | 4739 / 4733 | 0.00277 | 0.00225 / 0.00225 | 0.00227 / 0.00229 |
| *Pinaroloxias inornata* | Cocos | 8 | 4657 / 4654 | 0.00123 | 0.00121 / 0.00122 | 0.00120 / 0.00119 |
| *Ovis aries* | Oula | 10 | 4167 / 4149 | 0.00390 | 0.00288 / 0.00279 | 0.00275 / 0.00277 |
|  | Panou | 10 | 4162 / 4143 | 0.00357 | 0.00265 / 0.00270 | 0.00258 / 0.00266 |

**Figure 2.**
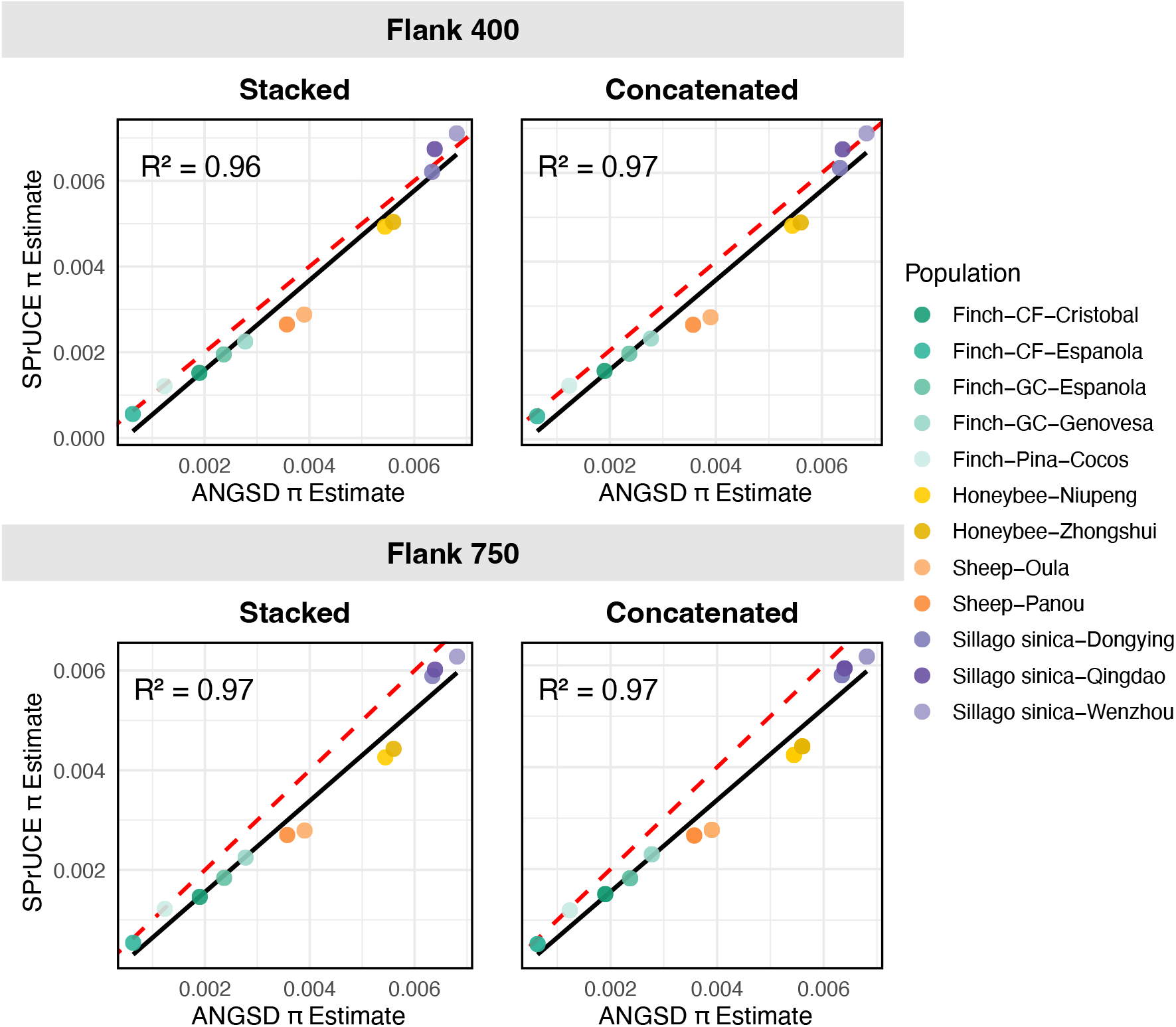
Comparisons of estimates of *π* from whole genome data using ANGSD versus SPrUCE UCE estimates for flank sizes of 400 and 750. Pearson’s correlation coefficient was used to assess agreement, and the coefficient of determination (R^2^) is reported for each comparison. The dashed red line represents the 1:1 relationship (x = y), and the solid black line shows the fitted linear regression. See also Table 1.

Beyond correlation, the *π* values estimated by SPrUCE are also close to ANGSD, with a tendency to estimate slightly lower values (11% on average). The only species where the differences are substantial is sheep, where SPrUCE’s estimates are 26% lower than ANGSD. Conversely, estimates are highly similar for the smelt dataset, which has the highest number of individuals, followed by the finch. In particular, SPrUCE is sensitive enough to detect diversity levels as low as 0.0005 for the finch species *Certhidea fusca*, and identified Cristobal as more diverse than Espanola, matching the same patterns as ANGSD. Estimates using 400bp flanks tend to be slightly higher than 750bp, especially in the Stacked mode (6% on average). The *π* estimates from the Stacked and Concatenated algorithms are very similar.

### 3.2 Comparing the Gompertz function to alternatives

Across all populations, the Gompertz-based estimates (SPrUCE) yielded estimates that were far closer to ANGSD than uncorrected mean *π* values (Fig. 3). Gompertz estimates are on average roughly 20% (0–42%) larger at 750bp flank and 50% (0–83%) at 400bp compared to uncorrected values, and thus, dramatically reduce the underestimation of *π* from UCEs compared to genome-wide estimates using ANGSD (Fig. S4). The only populations where Gompertz does not increase the estimate were *C. fusca* Espanola and *P. inornata* Cocos.

**Figure 3.**
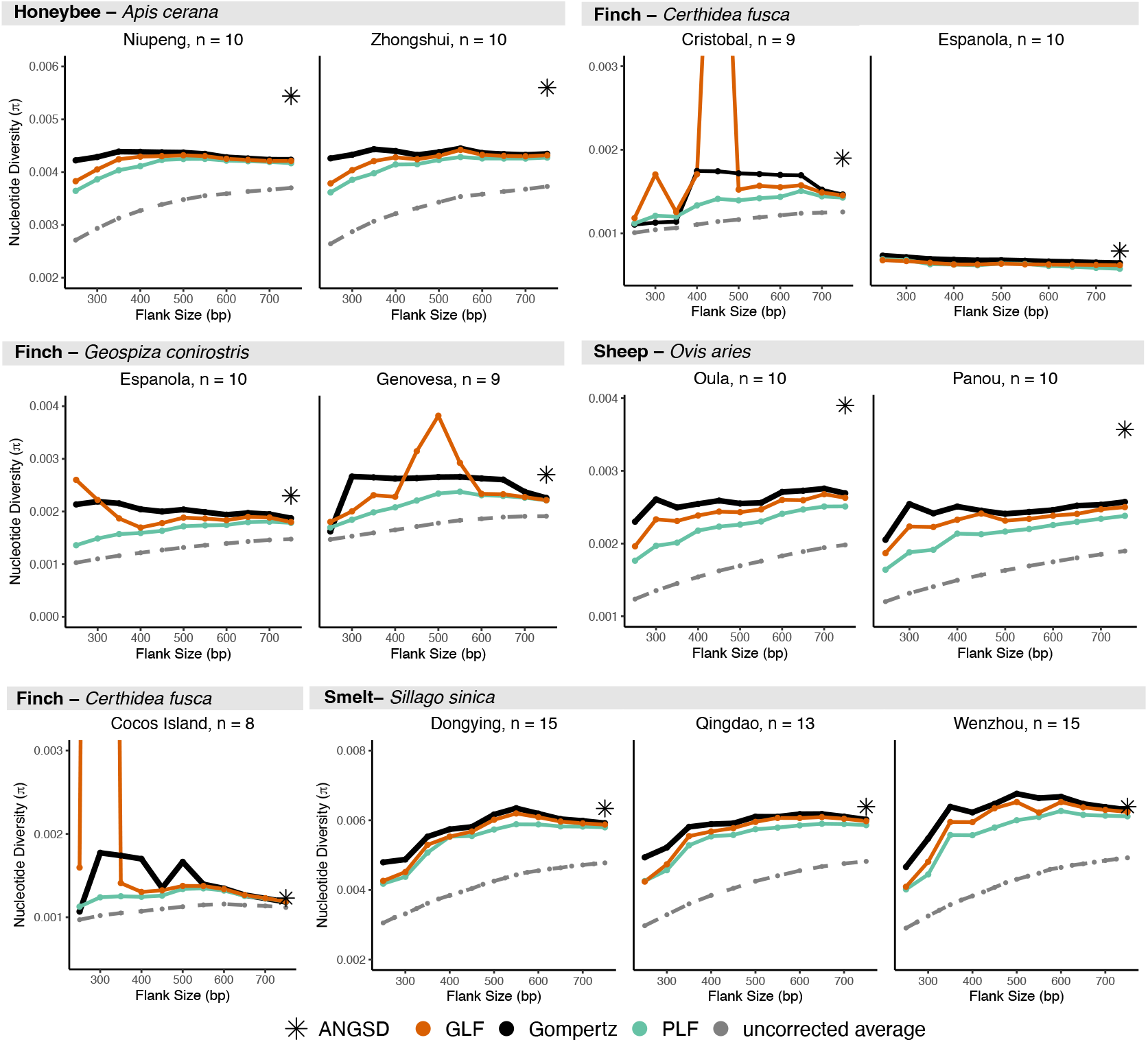
Gompertz Function fit compared to Generalized Logistic Function (GLF), and Piecewise Linear Function for decreasing flank sizes.

The advantage of the Gompertz function is further highlighted when testing robustness to flank size (Fig. 3). As the flanks become smaller, the gap between SPrUCE and uncorrected values further increases. This is because, unlike the uncorrected average, Gompertz estimates are relatively stable to the flank size used. In particular, they are largely consistent from 400 to 750 bp flanks and remained quite stable even for shorter flanks in *most* datasets. Only below 300 bp flank do we start to see dramatic reductions in the estimates, and even then, only for some species. Cases with 20% or more change in the *π* estimate compared to 750bp flank are observed only when flank sizes are 250bp, except for a single population (*Pinaroloxias inornata*), which showed less stability.

Besides Gompertz, we tested correcting UCE distances using the simpler PLF and the more parameter-rich GLF functions. The more parameterized GLF model exhibited instability and erratic behavior (spikes), especially in the fish and finch datasets (Fig. 3). Results show that as the flank size decreases, the six parameters of GLF become difficult to fit reliably with limited data, resulting in estimates that change widely. Neither Gompertz nor the simpler PLF suffered from such instability. PLF function produced relatively stable estimates close to Gompertz. However, PLF was less stable than Gompertz as the flank size shrank.

In summary, as we shrank the flank size, the 3-parameter Gompertz model was more robust than the 6-parameter GLF, more accurate than the 2-parameter PLF, and far more accurate than the uncorrected means. Estimates remain largely accurate for 300bp or longer flanks, but accuracy drops if even smaller flanks are used.

### 3.3 Impact of subsampling, individuals, and dataset completeness

For both 400 bp and 750 bp flanking regions, random subsampling of UCE loci showed that nucleotide diversity (*π*) estimates stabilized once more than 100 loci were included (Fig. 4a, S5a). Both Stacked and Concatenated modes produced estimates with low level of variation across replicates given enough loci, with only small improvements beyond 200 loci. For example, with Stacked and 750bp, the coefficient of variance of *π* estimates across replicates drops below 0.1 at 200 loci for all populations; at 800 loci, it ranges between 0.01 to 0.06.

**Figure 4.**
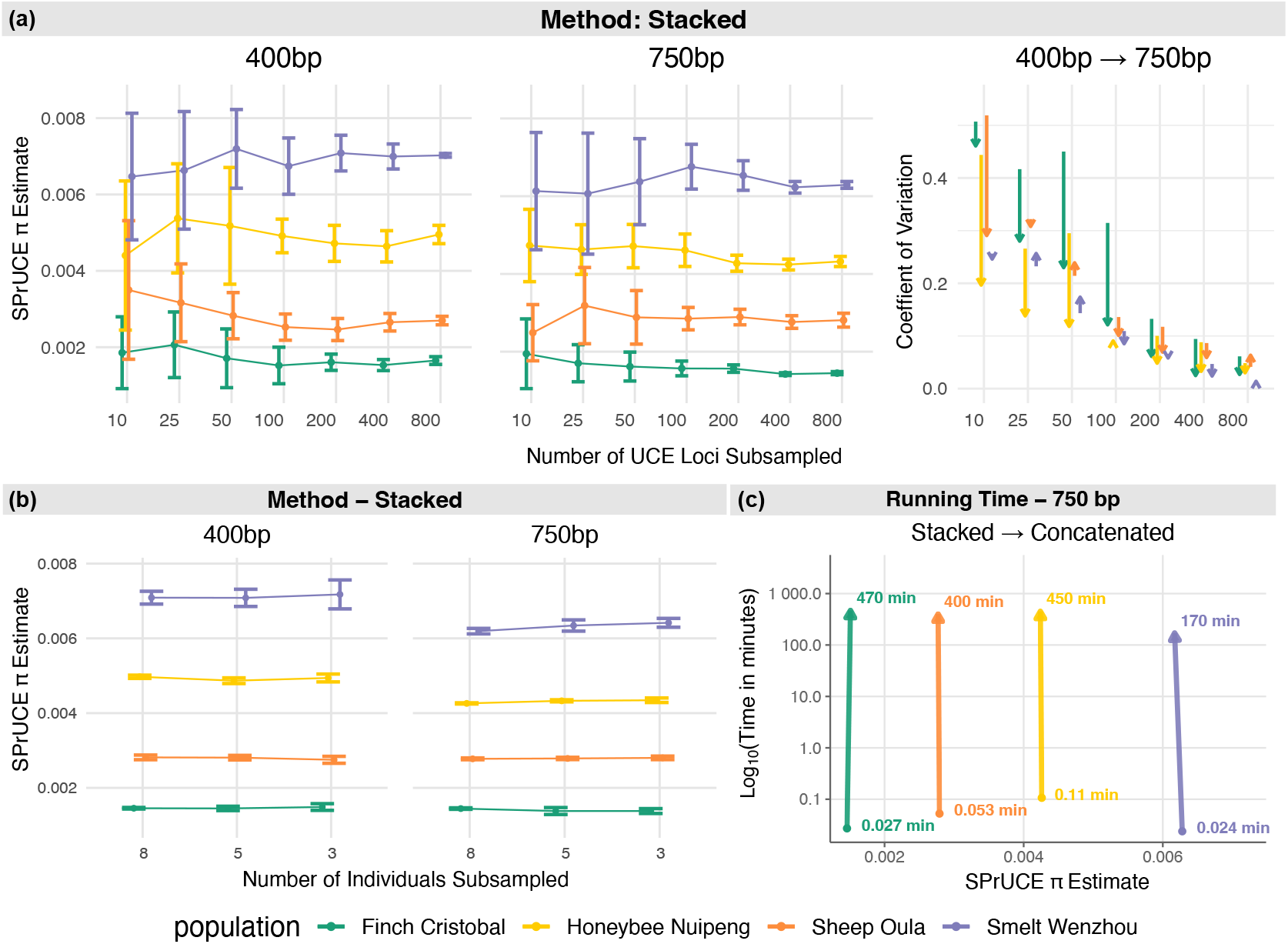
Estimates of nucleotide diversity (*π*) from random subsampling of UCE loci and individuals. (a) SPrUCE *π* estimates from subsampling UCE loci (10–800 loci, ten replicates each) using the Stacked mode at 400 bp and 750 bp flanking regions; coefficients of variation are shown at right. (b) Estimates obtained by varying the number of individuals per population (3, 5, and 8 individuals, ten replicates each) using the Stacked mode. (c) Runtime comparison between the Stacked and Concatenated modes at 750 bp flank length using a single CPU core. Population colors correspond to Finch (Green), Honeybee (Yellow), Sheep (Orange), and Smelt (Purple).

When comparing flank lengths, when the number of loci is small, the 400 bp analyses generally produced more variable estimates than the 750 bp analyses (Fig. 4a, S5a). The reduced variability of 750bp compared to 400bp is pronounced when 100 loci or fewer are available, but diminishes as the number of loci increases; at 800 loci, very little difference is observed between 400bp and 750bp. The use of 750bp versus 400 flanks makes a substantial difference in the variability for honeybee and finch datasets, but has a negligible impact on smelt. Across subsampling levels, the Concatenated mode exhibited slightly broader ranges compared to the Stacked mode in almost all datasets.

Subsampling individuals had minimal effect on SPrUCE nucleotide diversity (*π*) estimates across datasets (Fig. 4b). Estimates were highly consistent when reducing the sample size from 8 to 5 individuals, with only a slight increase in variance observed when subsampling to 3 individuals. The smelt dataset exhibited the greatest variability, particularly at 400 bp flanks under the Stacked mode; however, this effect was reduced at 750 bp and a similar pattern was observed in the Concatenated mode (Figs. 4b, S5b). The smelt dataset contained the fewest available UCE loci (898 or fewer; see Table S5), suggesting that the combined effects of the reduced number of loci and sample size alone can contribute to higher variability in estimates. Similar to subsampling the number of individuals, filtering loci by completeness threshold also had little to no effect on *π* estimates in either Stacked or Concatenated modes across both 400 bp and 750 bp flanking regions (Fig. S6).

### 3.4 Running time

Runtime comparisons revealed substantial differences between the two SPrUCE modes (Fig. 4c; Table S6). The Stacked mode was consistently faster across all datasets and flank lengths, completing in under 7 seconds for all populations (400 bp: 1.4–4.7 s; 750 bp: 1.4–6.4 s) when run on a single CPU core (Fig S7). In contrast, runtime for the Concatenated mode increased dramatically with dataset size, ranging from 30 minutes to 7 hours at 400 bp flanks and 1.5–13.5 hours at 750 bp flanks. On average, Stacked was around 5,000*×* faster than Concatenated. Despite the large runtime disparity, both methods produced highly consistent estimates of nucleotide diversity (*π*) across all datasets. The slowdown in the Concatenated mode results from independently handling large multi-locus alignments, which increases memory usage and computation time during matrix construction and model fitting.

### 3.5 Application to real UCE datasets

For the Willet dataset, SPrUCE estimates of nucleotide diversity (*π*) for the two subspecies of *Tringa semipalmata* were *π* = 0.00113 for the eastern subspecies (*T. s. semipalmata*), and *π* = 0.00170 for the western subspecies (*T. s. inornata*) for Stacked mode, with consistent estimates using Concatenated (Table 2). These diversity values translated to effective population size (*N*_*e*_) estimates of 185k – 195k for *T. s. semipalmata* and 278k – 314k for *T. s. inornata*. While *N*_*e*_ is not expected to always match the actual population size, these results were mostly congruent with estimates of the number of breeding-aged individuals (250k) for *Tringa semipalmata* (Partners in Flight, 2025). We note that using a phylogenetic analysis, Oswald et al. 2016 provided a much larger *N*_*e*_ estimate of 790k (*T. s. semipalmata*) and 2M (*T. s. inornata*), which far exceed the estimated number of individuals.

**Table 2:** Comparison of nucleotide diversity (*π*) estimates from published papers and SPrUCE analyses.

| Species | Sample Size<br>(n) | Paper<br>$\pi$ | SPrUCE<br>Stacked | SPrUCE<br>Concatenated |
| --- | --- | --- | --- | --- |
| *Plectrophenax nivalis* | $4 \times 2$ | 0.000523 | 0.000467 (64k) | 0.000526 (72k) |
| *Plectrophenax hyperboreus* | $4 \times 2$ | 0.000493 | 0.000398 (54k) | 0.000494 (67k) |
| *Tringa semipalmata inornata* | 19 | * | 0.00170 (278k) | 0.00192 (314k) |
| *Tringa semipalmata semipalmata* | 11 | * | 0.00113 (184k) | 0.00119 (194k) |

Results for the Winker et al. 2018 dataset of snow buntings (*Plectrophenax nivalis*) and McKay’s buntings (*Plectrophenax hyperboreus*) showed a close correspondence between our estimates and the published values, showing low levels of diversity (Table 2). We note that, unique among our datasets, the Concatenated mode resulted in somewhat higher (12%) estimates on this dataset compared to Stacked. The Concatenated estimates better match the estimates by Winker et al. 2018, indicating that they may be more accurate. We note that in these real datasets, UCEs have more length heterogeneity than the WGS-extracted UCEs. In addition, our estimates of effective population size (*N*_*e*_) followed expected trends for these species, with *P. nivalis* showing larger *N*_*e*_ (64k – 72k) relative to the more range-restricted *P. hyperboreus* (55k – 68k). For McKay buntings, which have a limited range and are well represented by the sampling of Winker et al. (2018), the estimated *N*_*e*_ was congruent with the number of breeding individuals estimated at 31k as indicated by Partners in Flight (2025). For snow bunting, the sampling is restricted to a small region of Alaska, whereas the global population has a much larger range, making it impossible to compare the estimated *N*_*e*_ and the census counts. The estimates of *N*_*e*_ by Winker et al. 2018 according to a split-migration model were about 2 – 3 *×* higher than our Stacked *N*_*e*_ estimates and 1.5 – 2.5*×* higher than Concatenated.

## 4 Discussion

There is a critical need for scalable approaches that can accurately capture genome-wide patterns of genetic diversity in non-model organisms, especially when working with low-coverage data. Full reference genomes remain unavailable or incomplete for many taxa, and whole-genome resequencing (WGS) pipelines are time-consuming and computationally demanding. Ultraconserved elements (UCEs) offer a powerful alternative: Once a bait set is designed, it can be applied broadly across taxa, often across entire families or orders (Faircloth et al., 2015). Moroever, they can be used to generate a SNP matrix without the need for a reference genome. These advantages make UCEs an increasingly attractive biomarker for genomic monitoring for conservation purposes. Our tool SPrUCE builds on this foundation by enabling rapid and accurate estimation of nucleotide diversity (*π*) directly from UCE alignments. Specifcally, it improves upon direct estimates by correcting for loss of diversity in ultraconserved regions.

Across four representative taxa (honeybee, finch, smelt, and sheep) and three UCE bait sets, SPrUCE produced *π* estimates that closely matched those from ANGSD, despite relying on very different computational paradigms (Korneliussen et al., 2014). ANGSD estimates nucleotide diversity using likelihood-based methods that require mapping reads to a reference genome, whereas SPrUCE operates reference-free by extracting allele frequencies directly from UCE alignments. The agreement between these two fundamentally different approaches reinforces the robustness of UCE data for estimating population-level genetic diversity. Another important difference lies in computational cost and input requirements: ANGSD may require hours to days of compute time per dataset (Table S7), depending on sample/genome size, and sequencing coverage. In contrast, SPrUCE runs on lightweight UCE alignments (e.g., MAFFT output from Phyluce) and can complete analyses in under 10 seconds per population using a single CPU core. This efficiency enables large-scale analyses and makes SPrUCE practical for settings with limited computational resources available. SPrUCE also offers practical advantages over other reduced-representation approaches. While perhaps sufficient for topological inference in phylogenetic studies (Manthey et al., 2016), RADseq suffers from allele dropout in highly diverged taxa and high levels of missing data, which can bias its diversity estimates (Arnold et al., 2013). In contrast, UCE capture works reliably on fragmented DNA. Mitochondrial DNA remains widely used in conservation genetics, but mtDNA has dynamics (such as mutation rates, maternal inheritance) that can significantly deviate from genome-wide patterns (Ferreira et al., 2024). By incorporating thousands of loci across the nuclear genome, SPrUCE provides a more comprehensive view of genetic diversity, while keeping the cost manageable.

Subsampling results revealed that longer flanking regions produced less variable estimates, suggesting that an extended flanking sequence (700bp+) provides additional informative polymorphisms. However, increasing the number of loci proved even more fruitful, as it impacted the variance in estimates more than other factors. While more loci are better, we observed that stable estimates of *π* could be achieved with as few as 400 UCE loci and five individuals per population, and are robust to missing data and incomplete loci. UCE bait sets may also differ in their genomic distribution across taxa, which can influence diversity estimates. For example, using the tetrapod bait set, birds showed strong agreement between ANGSD and SPrUCE estimates, while domesticated sheep exhibited slightly lower diversity values. This pattern may reflect how UCEs are organized across genomes in different lineages, potentially sampling regions with distinct evolutionary constraints or mutation rates. These bait-specific and taxon-specific biases warrant further evaluation and motivate future comparative work. Note that studies can generate their own curated bait sets tailored to specific clades, which is straightforward to implement using pipelines such as Phyluce (Faircloth, 2016) or Deduce (Cummins et al., 2024). However, in this study, we relied on publicly available UCE bait sets to emulate situations where ecologists use existing sets to minimize cost.

The SPrUCE estimates were slightly lower than those from ANGSD, especially for the two sheep populations. We note that according to the Gompertz parameter *β*, which is related to the recombination rate, these sheep (as well as *C. fusca Espanola*) populations stand out as having lower recombination rates (Fig. S8). A lower recombination rate can make it more difficult for Gompertz to find the correct asymptote given limited flanks, as evident from the fitted Gompertz functions across our datasets (Fig. S8). Thus, the lower values compared to ANGSD in these cases may mean that SPrUCE has some residual under-estimation. However, we note that ANGSD also provides higher estimates than the alternative methods that rely on SNP matrices instead of genotype likelihood (Korunes and Samuk, 2021). Thus, differences between ANGDS and SPrUCE could also be a result of over-estimation by ANGSD. We also compared our results to reference *π* estimates reported from each respective study (Table S6), which showed great concordance with SPrUCE for sheep (the only dataset where SPrUCE noticeably under-estimated ANGSD), reasonable similarity for finch, but more differences for honeybee or smelt. Because each study used different tools, filters, algorithms, variant calling thresholds, and diversity estimation pipelines, a direct comparison to these reported estimates is not feasible and beyond the scope of this study. These discrepancies likely reflect differences in filtering strategies and a host of other choices made in each whole-genome diversity study. Thus, without access to true values, it remains unclear whether the remaining small differences between ANGSD and SPrUCE stem from the inaccuracy of our model or particular choices of ANGSD parameters.

Benchmarking diversity estimators for UCE data is challenging because no simulation frameworks currently exist that replicate UCE architecture (conserved core and variable flanks) and their evolutionary properties. Existing coalescent simulators do not incorporate heterogeneity within and across UCE core into flanking regions, nor do they model probe design or capture bias, limiting their utility for method evaluation. Therefore, our validation of SPrUCE relied exclusively on empirical datasets, which required running ANGSD on full genome-aligned BAM files for comparison and substantially increased benchmarking runtime, and treating ANGSD as the standard for comparison. Developing UCE-aware simulation tools would be a valuable next step, enabling controlled assessments under known demographic and evolutionary scenarios.

Beyond evaluation, the method itself can be further improved in the future. We were able to theoretically justify the Gompertz function only for regimes with very low recombination rate and mutation rate compared to the selection coefficient. One could easily incorporate the recombination-aware corrections using the Hudson-Kaplan model (HK) shown earlier instead of Gompertz. We tried this approach and found that this HK-based correction was extremely sensitive to flank size; while its estimates at 750bp were as good as Gompertz, it immediately started to overestimate *π* as the flank size shrank (Fig. S9). These results suggest that UCEs may violate the equilibrium assumptions used by Hudson and Kaplan (1995) to derive their model. More realistic models need to be developed in the future. Also left to future work is extending SPrUCE beyond nucleotide diversity (*π*) to also estimate population differentiation metrics such as *F*_*ST*_.

Between the two SPrUCE modes, Stacked seems sufficient for most uses. Computational runtime across datasets showed that the Stacked is approximately 5,000*×* faster than the Concatenated mode, while producing nearly identical estimates of *π* (Table S6). Stacked aggregates diversity estimates across all loci, making it computationally efficient. Theory suggests Concatenated may be more robust to errors in the data, extreme length heterogeneity, or lack of coverage, making it suitable only when such issues are suspected. For example, on the real snow bunting dataset with a substantial length heterogeneity, Concatenated estimates diverged from Stacked and were more consistent with results from Winker et al. (2018). Users who can afford to run both modes are encouraged to do so to compare results. Finally, SPrUCE is designed to be user-friendly, offering flexibility through multiple customizable filters, including flank size selection, choice of estimation method (-stacked and -concatenated), minimum base threshold for locus inclusion, and input format.

Overall, SPrUCE provides a flexible and efficient tool for quantifying genetic diversity from UCE data across a wide range of taxa. It reduces computational barriers and eliminates the need for reference genomes. SPrUCE makes large-scale genomic monitoring more accessible and opens new opportunities for biodiversity research and conservation genetics.

## 5 ACKNOWLEDGEMENTS

We thank the Minderoo Foundation for supporting portions of this work and providing access to genomic resources. We also thank the authors of the datasets used for genetic diversity comparisons, whose publicly available data made this study possible. We are especially grateful to Brant Faircloth for pioneering the development of UCE approaches and for inspiring continued innovation in UCE-based evolutionary genomics.

## 6 Author Contributions

Conception: VB, SM. Design: DM, AOBS, VB, SM. Implementation: AOBS, DM. Analysis: DM, VB, SM. Writing: All authors.

## 7 Code and Data Availability

Full details for pre-processing scripts procedures for reproducing data are available at: https://github.com/dairabel92/spruce_analysis. The SPrUCE software is publicly available at https://github.com/dairabel92/spruce.

