## Supplementary Material for "SPrUCE: Utilizing Ultraconserved Elements of DNA for Population-Level Genetic Diversity Estimation"

#### Contents

**S1 Supplementary Tables** 3

**S2 Supplementary Figures** 9

**S3 Supplementary text** 17
S3.1 SPrUCE Algorithm . . . 17
S3.2 Approximation of $\pi_i$ by $2f_i$ . . . 18

### S1 Supplementary Tables

| ID | Species | Population | NCBI ID | Genome Size | UCEs 400bp | UCEs 750bp |
|---|---|---|---|---|---|---|
| DY1 | *Sillago sinica* | Dongying | SRR23814189 | 557 Mb | 993 | 953 |
| DY2 | *Sillago sinica* | Dongying | SRR23814188 | 552 Mb | 992 | 952 |
| DY3 | *Sillago sinica* | Dongying | SRR23814174 | 551 Mb | 993 | 953 |
| DY4 | *Sillago sinica* | Dongying | SRR23814163 | 549 Mb | 990 | 953 |
| DY5 | *Sillago sinica* | Dongying | SRR23814152 | 548 Mb | 998 | 958 |
| DY6 | *Sillago sinica* | Dongying | SRR23814148 | 543 Mb | 997 | 962 |
| DY7 | *Sillago sinica* | Dongying | SRR23814147 | 548 Mb | 992 | 954 |
| DY8 | *Sillago sinica* | Dongying | SRR23814185 | 546 Mb | 997 | 961 |
| DY9 | *Sillago sinica* | Dongying | SRR23814184 | 557 Mb | 995 | 955 |
| DY10 | *Sillago sinica* | Dongying | SRR23814182 | 527 Mb | 990 | 963 |
| DY11 | *Sillago sinica* | Dongying | SRR23814187 | 544 Mb | 993 | 957 |
| DY12 | *Sillago sinica* | Dongying | SRR23814186 | 545 Mb | 995 | 959 |
| DY13 | *Sillago sinica* | Dongying | SRR23814183 | 546 Mb | 992 | 956 |
| DY14 | *Sillago sinica* | Dongying | SRR23814181 | 547 Mb | 989 | 948 |
| DY15 | *Sillago sinica* | Dongying | SRR23814180 | 546 Mb | 994 | 956 |
| QD1 | *Sillago sinica* | Qingdao | SRR23814179 | 549 Mb | 987 | 948 |
| QD2 | *Sillago sinica* | Qingdao | SRR23814178 | 552 Mb | 991 | 953 |
| QD3 | *Sillago sinica* | Qingdao | SRR23814177 | 555 Mb | 990 | 950 |
| QD4 | *Sillago sinica* | Qingdao | SRR23814176 | 556 Mb | 991 | 956 |
| QD5 | *Sillago sinica* | Qingdao | SRR23814175 | 549 Mb | 994 | 956 |
| QD6 | *Sillago sinica* | Qingdao | SRR23814173 | 550 Mb | 993 | 956 |
| QD7 | *Sillago sinica* | Qingdao | SRR23814172 | 544 Mb | 994 | 957 |
| QD8 | *Sillago sinica* | Qingdao | SRR23814171 | 550 Mb | 985 | 948 |
| QD9 | *Sillago sinica* | Qingdao | SRR23814170 | 554 Mb | 991 | 953 |
| QD10 | *Sillago sinica* | Qingdao | SRR23814169 | 544 Mb | 993 | 957 |
| QD11 | *Sillago sinica* | Qingdao | SRR23814168 | 542 Mb | 987 | 947 |
| QD12 | *Sillago sinica* | Qingdao | SRR23814167 | 538 Mb | 990 | 951 |
| QD13 | *Sillago sinica* | Qingdao | SRR23814166 | 544 Mb | 991 | 952 |
| WZ1 | *Sillago sinica* | Wenzhou | SRR23814165 | 550 Mb | 987 | 950 |
| WZ2 | *Sillago sinica* | Wenzhou | SRR23814164 | 552 Mb | 996 | 957 |
| WZ3 | *Sillago sinica* | Wenzhou | SRR23814162 | 557 Mb | 983 | 942 |
| WZ4 | *Sillago sinica* | Wenzhou | SRR23814161 | 551 Mb | 982 | 943 |
| WZ5 | *Sillago sinica* | Wenzhou | SRR23814160 | 553 Mb | 991 | 954 |
| WZ6 | *Sillago sinica* | Wenzhou | SRR23814159 | 552 Mb | 986 | 947 |
| WZ7 | *Sillago sinica* | Wenzhou | SRR23814158 | 553 Mb | 987 | 946 |
| WZ8 | *Sillago sinica* | Wenzhou | SRR23814157 | 547 Mb | 990 | 949 |
| WZ9 | *Sillago sinica* | Wenzhou | SRR23814156 | 558 Mb | 992 | 953 |
| WZ10 | *Sillago sinica* | Wenzhou | SRR23814155 | 553 Mb | 989 | 949 |
| WZ11 | *Sillago sinica* | Wenzhou | SRR23814154 | 544 Mb | 992 | 954 |
| WZ12 | *Sillago sinica* | Wenzhou | SRR23814153 | 550 Mb | 993 | 956 |
| WZ13 | *Sillago sinica* | Wenzhou | SRR23814151 | 550 Mb | 997 | 956 |
| WZ14 | *Sillago sinica* | Wenzhou | SRR23814150 | 549 Mb | 981 | 945 |
| WZ15 | *Sillago sinica* | Wenzhou | SRR23814149 | 547 Mb | 996 | 958 |

Table S1: Sample metadata from Zhao et al. 2023 for *Sillago sinica* populations in China (Dongying, Qingdao, and Wenzhou), including ID for this study, species, population, SRR ID, MEGAHIT genome size after assembly, and number of loci recovered with 400 bp and 750 bp flanking regions. Raw data come from project PRJNA936440.

| ID | Species | Population | NCBI ID | Genome Size | UCEs 400bp | UCEs 750bp |
|---|---|---|---|---|---|---|
| cfE1 | *Certhidea fusca* | Espanola | SRR1607346 | 908 Mb | 3890 | 3884 |
| cfE2 | *Certhidea fusca* | Espanola | SRR1607350 | 948 Mb | 4265 | 4257 |
| cfE3 | *Certhidea fusca* | Espanola | SRR1607351 | 882 Mb | 4062 | 4049 |
| cfE4 | *Certhidea fusca* | Espanola | SRR1607354 | 914 Mb | 3990 | 3979 |
| cfE5 | *Certhidea fusca* | Espanola | SRR1607356 | 878 Mb | 3919 | 3906 |
| cfE6 | *Certhidea fusca* | Espanola | SRR1607357 | 940 Mb | 4405 | 4390 |
| cfE7 | *Certhidea fusca* | Espanola | SRR1607359 | 999 Mb | 4536 | 4525 |
| cfE8 | *Certhidea fusca* | Espanola | SRR1607361 | 952 Mb | 4276 | 4271 |
| cfE9 | *Certhidea fusca* | Espanola | SRR1607363 | 923 Mb | 4102 | 4093 |
| cfE10 | *Certhidea fusca* | Espanola | SRR1607348 | 921 Mb | 4042 | 4031 |
| cfC1 | *Certhidea fusca* | Cristobal | SRR1607326 | 952 Mb | 4297 | 4288 |
| cfC2 | *Certhidea fusca* | Cristobal | SRR1607330 | 911 Mb | 3960 | 3951 |
| cfC3 | *Certhidea fusca* | Cristobal | SRR1607332 | 923 Mb | 4187 | 4179 |
| cfC4 | *Certhidea fusca* | Cristobal | SRR1607334 | 912 Mb | 4168 | 4095 |
| cfC5 | *Certhidea fusca* | Cristobal | SRR1607336 | 957 Mb | 4311 | 4302 |
| cfC6 | *Certhidea fusca* | Cristobal | SRR1607337 | 959 Mb | 4278 | 4272 |
| cfC8 | *Certhidea fusca* | Cristobal | SRR1607342 | 864 Mb | 3881 | 3873 |
| cfC9 | *Certhidea fusca* | Cristobal | SRR1607343 | 830 Mb | 3671 | 3661 |
| cfC10 | *Certhidea fusca* | Cristobal | SRR1607327 | 979 Mb | 4478 | 4467 |
| gcE1 | *Geospiza conirostris* | Espanola | SRR1607296 | 922 Mb | 4106 | 4095 |
| gcE2 | *Geospiza conirostris* | Espanola | SRR1607302 | 969 Mb | 4417 | 4407 |
| gcE3 | *Geospiza conirostris* | Espanola | SRR1607306 | 1014 Mb | 4619 | 4604 |
| gcE4 | *Geospiza conirostris* | Espanola | SRR1607310 | 960 Mb | 4394 | 4382 |
| gcE5 | *Geospiza conirostris* | Espanola | SRR1607312 | 977 Mb | 4447 | 4436 |
| gcE6 | *Geospiza conirostris* | Espanola | SRR1607316 | 991 Mb | 4543 | 4531 |
| gcE7 | *Geospiza conirostris* | Espanola | SRR1607318 | 931 Mb | 4203 | 4191 |
| gcE8 | *Geospiza conirostris* | Espanola | SRR1607320 | 991 Mb | 4513 | 4500 |
| gcE9 | *Geospiza conirostris* | Espanola | SRR1607323 | 995 Mb | 4571 | 4561 |
| gcE10 | *Geospiza conirostris* | Espanola | SRR1607299 | 938 Mb | 4240 | 4229 |
| gcG1 | *Geospiza conirostris* | Genovesa | SRR1607365 | 962 Mb | 4347 | 4343 |
| gcG2 | *Geospiza conirostris* | Genovesa | SRR1607369 | 968 Mb | 4376 | 4364 |
| gcG3 | *Geospiza conirostris* | Genovesa | SRR1607372 | 881 Mb | 3981 | 3977 |
| gcG4 | *Geospiza conirostris* | Genovesa | SRR1607373 | 992 Mb | 4579 | 4563 |
| gcG5 | *Geospiza conirostris* | Genovesa | SRR1607375 | 913 Mb | 4037 | 4027 |
| gcG6 | *Geospiza conirostris* | Genovesa | SRR1607377 | 873 Mb | 4019 | 4009 |
| gcG7 | *Geospiza conirostris* | Genovesa | SRR1607380 | 878 Mb | 3755 | 3753 |
| gcG8 | *Geospiza conirostris* | Genovesa | SRR1607382 | 888 Mb | 3876 | 3872 |
| gcG9 | *Geospiza conirostris* | Genovesa | SRR1607384 | 898 Mb | 3899 | 3895 |
| pnC1 | *Pinaroloxias inornata* | Cocos Island | SRR1607513 | 764 Mb | 3399 | 3395 |
| pnC2 | *Pinaroloxias inornata* | Cocos Island | SRR1607514 | 908 Mb | 4031 | 4028 |
| pnC3 | *Pinaroloxias inornata* | Cocos Island | SRR1607517 | 817 Mb | 3663 | 3660 |
| pnC4 | *Pinaroloxias inornata* | Cocos Island | SRR1607518 | 843 Mb | 3704 | 3699 |
| pnC5 | *Pinaroloxias inornata* | Cocos Island | SRR1607520 | 842 Mb | 3604 | 3595 |
| pnC6 | *Pinaroloxias inornata* | Cocos Island | SRR1607522 | 763 Mb | 3259 | 3257 |
| pnC7 | *Pinaroloxias inornata* | Cocos Island | SRR1607528 | 996 Mb | 4566 | 4554 |
| pnC8 | *Pinaroloxias inornata* | Cocos Island | SRR1607531 | 894 Mb | 4022 | 4011 |

Table S2: Sample metadata from Lamichhaney et al. (2015) for finch species, including ID for this study, species, population, NCBI ID, genome size after assembly, and number of loci recovered with 400 bp and 750 bp flanking regions. Raw data comes from project PRJNA263122.

| ID | Species | Population | NCBI ID | Genome Size | UCEs 400bp | UCEs 750bp |
|---|---|---|---|---|---|---|
| NP1 | *Apis cerana* | Niupeng | SRR27281405 | 214 Mb | 2305 | 2297 |
| NP2 | *Apis cerana* | Niupeng | SRR27281404 | 210 Mb | 2343 | 2336 |
| NP3 | *Apis cerana* | Niupeng | SRR27281403 | 206 Mb | 2348 | 2345 |
| NP4 | *Apis cerana* | Niupeng | SRR27281402 | 208 Mb | 2366 | 2361 |
| NP5 | *Apis cerana* | Niupeng | SRR27281514 | 211 Mb | 2367 | 2362 |
| NP6 | *Apis cerana* | Niupeng | SRR27281513 | 206 Mb | 2333 | 2326 |
| NP7 | *Apis cerana* | Niupeng | SRR27281512 | 212 Mb | 2374 | 2366 |
| NP8 | *Apis cerana* | Niupeng | SRR27281511 | 205 Mb | 2338 | 2329 |
| NP9 | *Apis cerana* | Niupeng | SRR27281510 | 207 Mb | 2348 | 2341 |
| NP10 | *Apis cerana* | Niupeng | SRR27281509 | 205 Mb | 2309 | 2303 |
| ZS1 | *Apis cerana* | Zhongshui | SRR27281517 | 202 Mb | 2334 | 2328 |
| ZS2 | *Apis cerana* | Zhongshui | SRR27281516 | 203 Mb | 2349 | 2341 |
| ZS3 | *Apis cerana* | Zhongshui | SRR27281489 | 203 Mb | 2326 | 2319 |
| ZS4 | *Apis cerana* | Zhongshui | SRR27281446 | 205 Mb | 2368 | 2360 |
| ZS5 | *Apis cerana* | Zhongshui | SRR27281435 | 196 Mb | 2332 | 2322 |
| ZS6 | *Apis cerana* | Zhongshui | SRR27281424 | 206 Mb | 2352 | 2345 |
| ZS7 | *Apis cerana* | Zhongshui | SRR27281477 | 208 Mb | 2361 | 2353 |
| ZS8 | *Apis cerana* | Zhongshui | SRR27281466 | 201 Mb | 2316 | 2306 |
| ZS9 | *Apis cerana* | Zhongshui | SRR27281455 | 190 Mb | 2306 | 2296 |
| ZS10 | *Apis cerana* | Zhongshui | SRR27281412 | 194 Mb | 2360 | 2353 |

Table S3: Sample metadata from Wang et al. (2024) for *Apis cerana* individuals from Niupeng and Zhongshui populations, including ID for this study, species, population, NCBI ID, genome size after assembly, and number of loci recovered with 400 bp and 750 bp flanking regions. Raw data comes from project PRJNA1054499

| ID | Species | Population | NCBI ID | Genome Size | UCEs 400bp | UCEs 750bp |
|---|---|---|---|---|---|---|
| OL-1 | *Ovis aries* | Oula | SRR17643011 | 2.75 G | 4070 | 4050 |
| OL-2 | *Ovis aries* | Oula | SRR17643010 | 2.66 G | 4077 | 4057 |
| OL-3 | *Ovis aries* | Oula | SRR17642999 | 2.64 G | 4045 | 4024 |
| OL-4 | *Ovis aries* | Oula | SRR17642998 | 2.67 G | 4026 | 4016 |
| OL-5 | *Ovis aries* | Oula | SRR17642997 | 2.60 G | 4005 | 3986 |
| OL-6 | *Ovis aries* | Oula | SRR17642996 | 2.62 G | 4050 | 4033 |
| OL-7 | *Ovis aries* | Oula | SRR17642995 | 2.65 G | 4090 | 4066 |
| OL-8 | *Ovis aries* | Oula | SRR17642994 | 2.67 G | 4106 | 4085 |
| OL-9 | *Ovis aries* | Oula | SRR17642993 | 2.54 G | 4112 | 4094 |
| OL-10 | *Ovis aries* | Oula | SRR17642992 | 2.59 G | 4117 | 4096 |
| PO-1 | *Ovis aries* | Panou | SRR17643009 | 2.62 G | 4115 | 4094 |
| PO-2 | *Ovis aries* | Panou | SRR17643008 | 2.63 G | 4076 | 4057 |
| PO-3 | *Ovis aries* | Panou | SRR17643007 | 2.64 G | 4062 | 4045 |
| PO-4 | *Ovis aries* | Panou | SRR17643006 | 2.58 G | 4103 | 4082 |
| PO-5 | *Ovis aries* | Panou | SRR17643005 | 2.57 G | 4008 | 3989 |
| PO-6 | *Ovis aries* | Panou | SRR17643004 | 2.63 G | 4034 | 4022 |
| PO-7 | *Ovis aries* | Panou | SRR17643003 | 2.67 G | 4040 | 4018 |
| PO-8 | *Ovis aries* | Panou | SRR17643002 | 2.70 G | 4082 | 4057 |
| PO-9 | *Ovis aries* | Panou | SRR17643001 | 2.70 G | 3983 | 3959 |
| PO-10 | *Ovis aries* | Panou | SRR17643000 | 2.70 G | 4031 | 4009 |

Table S4: Sample metadata from Shi et al. (2023) for *Ovis aries* individuals from Oula and Panou populations, including ID for this study, species, population, NCBI ID, genome size after assembly, and number of loci recovered with 400 bp and 750 bp flanking regions. Raw data comes from project PRJNA797957

| Species Group | Population | Incomplete 400 / 750 | 50% Complete 400 / 750 | 80% Complete 400 / 750 | 100% Complete 400 / 750 |
|---|---|---|---|---|---|
| *A. cerana* | Niupeng | 2412 / 2409 | 2389 / 2382 | 2349 / 2340 | 1973 / 1965 |
| | Zhongshui | 2410 / 2402 | 2382 / 2373 | 2336 / 2326 | 1997 / 1987 |
| *C. fusca* | Cristobal | 4737 / 4729 | 4684 / 4674 | 4068 / 4053 | 1713 / 1706 |
| | Espanola | 4751 / 4744 | 4651 / 4635 | 3993 / 3477 | 1601 / 1597 |
| *G. conirostris* | Espanola | 4739 / 4728 | 4678 / 4662 | 4433 / 4417 | 2665 / 2653 |
| | Genovesa | 4739 / 4733 | 4655 / 4654 | 3933 / 3969 | 1694 / 1688 |
| *P. inornata* | Cocos Island | 4657 / 4654 | 4485 / 4476 | 3639 / 3625 | 1030 / 1029 |
| *O. aries* | Oula | 4167 / 4149 | 4148 / 4126 | 4096 / 4075 | 3496 / 3423 |
| | Panou | 4162 / 4143 | 4147 / 4125 | 4076 / 4051 | 3334 / 3314 |
| *S. sinica* | Dongying | 1005 / 971 | 1000 / 964 | 993 / 952 | 919 / 878 |
| | Qingdao | 1005 / 970 | 998 / 961 | 992 / 951 | 909 / 870 |
| | Wenzhou | 1005 / 970 | 999 / 958 | 988 / 947 | 898 / 861 |

Table S5: UCE loci recovered at different completeness thresholds (Incomplete, 50%, 80%, and 100%) for 400 bp and 750 bp flanks.

| Species Group | Population | STACKED | | CONCATENATED | |
|---|---|---|---|---|---|
| | | 400bp / Time(s) | 750bp / Time(s) | 400 bp/Time(s) | 750bp / Time(s) |
| *A. cerana* | Niupeng | 0.00493 / 4.7 | 0.00426 / 6.41 | 0.00481 / 15649.53 | 0.00424 / 27289.39 |
| | Zhongshui | 0.00504 / 3.22 | 0.00443 / 4.21 | 0.00488 / 16193.87 | 0.00441 / 22950.23 |
| *S. sinica* | Dongying | 0.00621 / 1.40 | 0.00589 / 3.22 | 0.00611 / 1725.15 | 0.00580 / 5093.64 |
| | Qingdao | 0.00674 / 2.62 | 0.00602 / 4.62 | 0.00653 / 5481.33 | 0.00594 / 5560.10 |
| | Wenzhou | 0.00711 / 3.95 | 0.00628 / 1.43 | 0.00689 / 3240.16 | 0.00617 / 10370.19 |
| *C. fusca* | Cristobal | 0.00152 / 2.17 | 0.00146 / 1.63 | 0.00154 / 15244.47 | 0.00151 / 28394.42 |
| | Espanola | 0.00056 / 2.62 | 0.00054 / 2.22 | 0.00052 / 5794.33 | 0.00053 / 8125.45 |
| *G. conirostris* | Espanola | 0.00195 / 2.13 | 0.00184 / 3.63 | 0.00193 / 14779.95 | 0.00182 / 48939.41 |
| | Genovesa | 0.00225 / 2.67 | 0.00225 / 2.98 | 0.00227 / 21278.82 | 0.00229 / 34843.64 |
| *P. inornata* | Cocos | 0.00121 / 3.03 | 0.00122 / 2.22 | 0.00121 / 9816.39 | 0.00119 / 14800.04 |
| *O. aries* | Oula | 0.00288 / 3.25 | 0.00279 / 3.16 | 0.00275 / 16673.69 | 0.00277 / 24245.68 |
| | Panou | 0.00265 / 2.51 | 0.00270 / 5.21 | 0.00258 / 25657.63 | 0.00266 / 21798.80 |

| Species | Pop. | ANGSD $\pi$ | Ref. Paper $\pi$ | Reference |
|---|---|---|---|---|
| *A. cerana* | Niupeng | 0.00544 | 0.00157 | Wang et al. 2024 |
| | Zhongshui | 0.00559 | 0.00158 | Wang et al. 2024 |
| *S. sinica* | Dongying | 0.00634 | 0.00241 | Zhao et al. 2023 |
| | Qingdao | 0.00639 | 0.00241 | Zhao et al. 2023 |
| | Wenzhou | 0.00681 | 0.00244 | Zhao et al. 2023 |
| *C. fusca* | Cristobal | 0.00190 | 0.0013 | Lamichhaney et al. 2015 |
| | Espanola | 0.00062 | 0.0003 | Lamichhaney et al. 2015 |
| *G. conirostris* | Espanola | 0.00236 | 0.0015 | Lamichhaney et al. 2015 |
| | Genovesa | 0.00277 | 0.0018 | Lamichhaney et al. 2015 |
| *P. inornata* | Cocos | 0.00123 | 0.0007 | Lamichhaney et al. 2015 |
| *O. aries* | Oula | 0.0039 | 0.00275 | Shit et al. 2023 |
| | Panou | 0.00357 | 0.00265 | Shit et al. 2023 |

Table S6: Top: Nucleotide diversity ($\pi$) and runtime (seconds) for UCE analyses using Stacked and Concatenated methods at 400 bp and 750 bp flanking regions. Bottom: The $\pi$ estimates from ANGSD and reference studies.

| Species | Common name | Population | CPU time (s) | Walltime (s) | $T_p$ |
|---|---|---|---|---|---|
| *Apis cerana* | Honeybee | Niupeng | 8837.23 | 55974 | 0.00544 |
| *Apis cerana* | Honeybee | Zhonshui | 8535.11 | 79778 | 0.00560 |
| *Sillago sinica* | Smelt | Dongying | 48188.80 | 201125 | 0.00634 |
| *Sillago sinica* | Smelt | Qingdao | 44783.30 | 185689 | 0.00639 |
| *Sillago sinica* | Smelt | Wenzhou | 52448.80 | 196087 | 0.00681 |
| *Certhidea fusca* | Finch | Cristobal | 30629.70 | 454237 | 0.00190 |
| *Certhidea fusca* | Finch | Española | 27450.97 | 380453 | 0.00063 |
| *Geospiza conirostris* | Finch | Española | 37672.08 | 261712 | 0.00237 |
| *Geospiza conirostris* | Finch | Genovesa | 33847.33 | 3057805 | 0.00277 |
| *Pinaroloxias inornata* | Finch | Cocos | 28902.48 | 292567 | 0.00123 |
| *Ovis aries* | Sheep | Oula | 103380.32 | 1047668 | 0.00390 |
| *Ovis aries* | Sheep | Panou | 82163.04 | 1049469 | 0.00357 |

Table S7: ANGSD computational time for `angsd -saf` and nucleotide diversity estimates ($T_p$).

### S2 Supplementary Figures

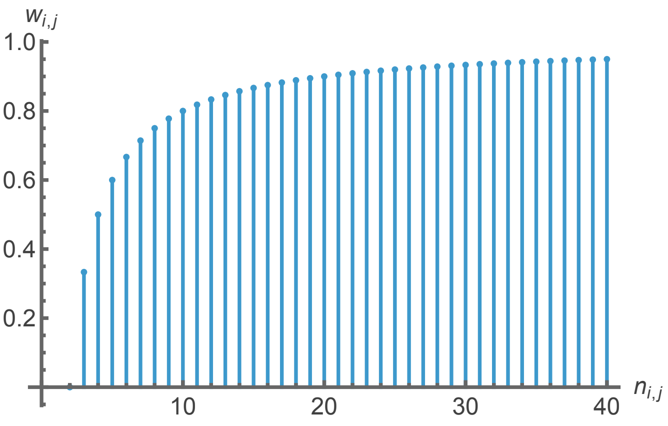

Figure S1: The weights of each site are proportional to $1 - \frac{2}{n+1}$, where $n$ is the number of elements present at the site. This function shows how weights grow and saturate for $n > 10$. Note that for $n = 1$, the weight is zero, as it should be, and as $n \to \infty$, weights go towards 1.

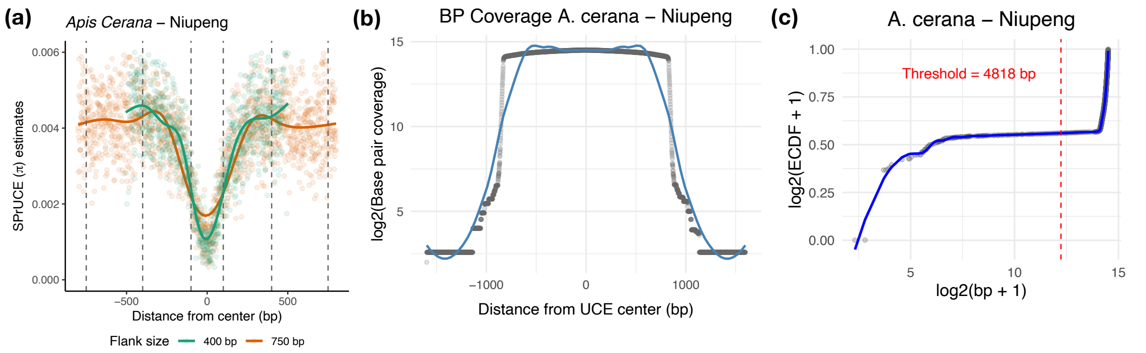

Figure S2: **(a)** UCE smilogram for 400 and 750 flank sizes for for *Apis cerana* (Honeybee – Niupeng). **(b)** Distribution of base pair coverage across positions flanking UCEs. Each point represents the log-transformed number of base pairs observed at a given position relative to the UCE center (position 0). A smoothed curve is overlaid to highlight the overall trend. Coverage is highest near the center of the UCE and decreases symmetrically toward the flanking regions. **(c)** Log-transformed spline-smoothed empirical cumulative distribution functions (ECDFs) of base pair counts. The dashed red line indicates the estimated inflection point based on the second derivative of the smoothed curve, used to define the base pair threshold for inclusion. Threshold values for bp included.

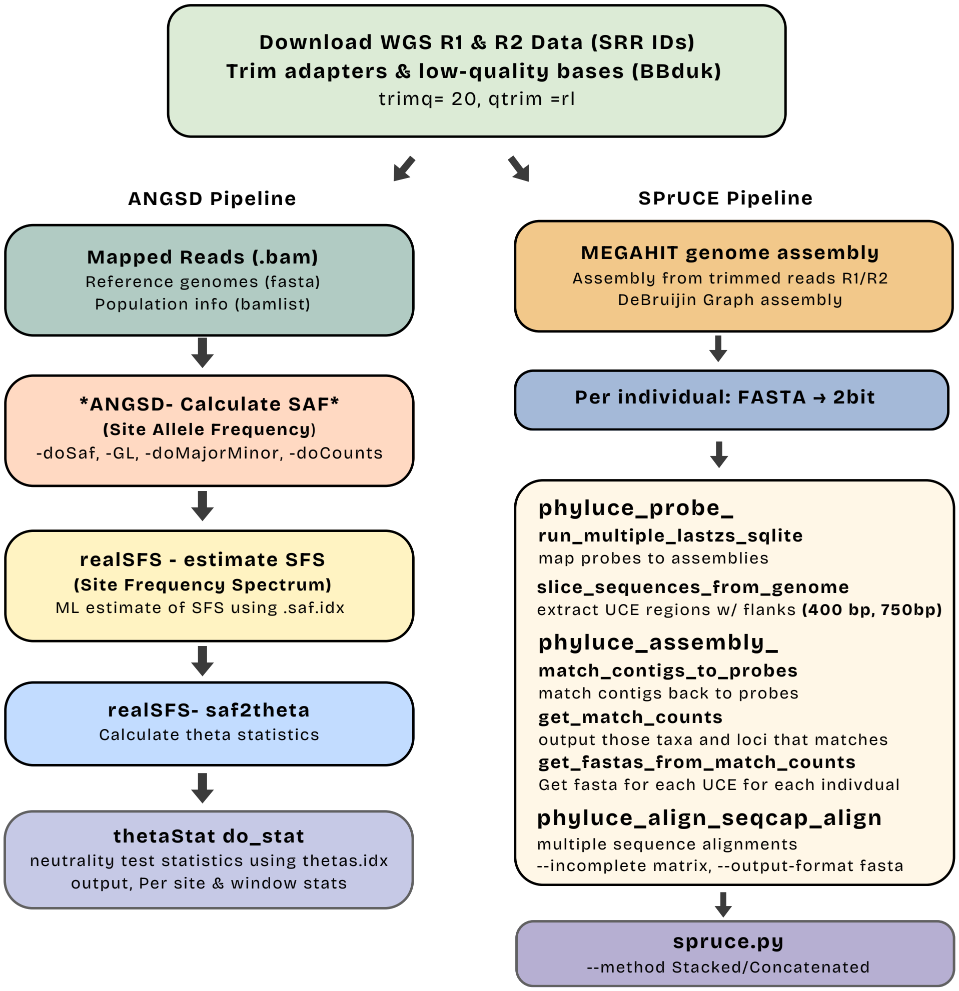

Figure S3: WGS data pre-processing pipeline for ANGSD and SPrUCE analysis

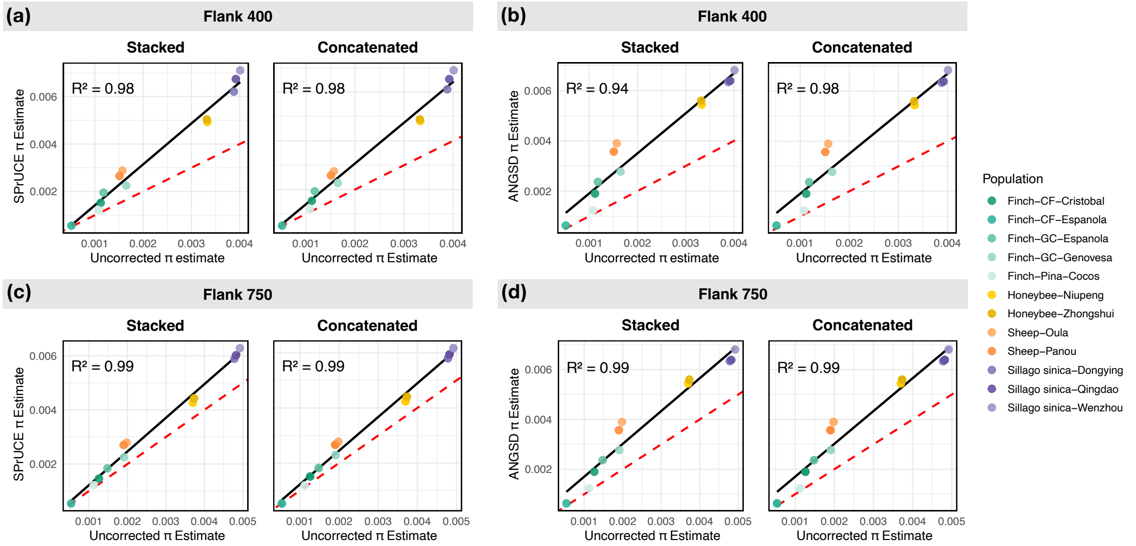

Figure S4: Comparison of uncorrected nucleotide diversity estimates with SPrUCE (left: a, c) and ANGSD (right: b, d) for 400 bp and 750 bp flanking regions. Pearson's correlation coefficient was used to assess agreement, and the coefficient of determination ($R^2$) is reported for each comparison. The dashed red line represents the 1:1 relationship ($x = y$), and the solid black line shows the fitted linear regression.

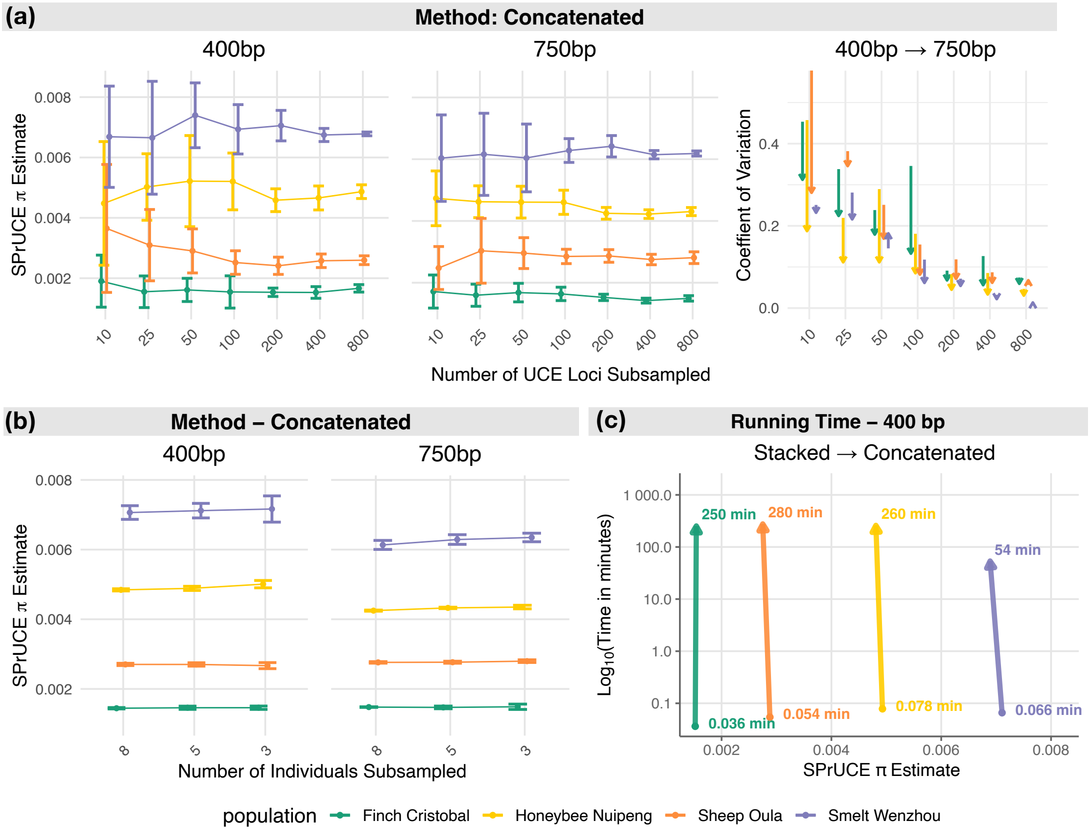

Figure S5: Estimates of nucleotide diversity $\pi$ from random subsampling of UCE loci and individuals. (a) SPrUCE $\pi$ estimates from subsampling UCE loci (10–800 loci, ten replicates each) using the Concatenated mode at 400 bp and 750 bp flanking regions; coefficients of variation are shown at right. (b) $\pi$ estimates obtained by varying the number of individuals per population (3, 5, and 8 individuals, ten replicates each) using the Concatenated mode. (c) Runtime comparison between the Stacked and Concatenated modes at 400 bp flank length using a single CPU core. Population colors correspond to Finch (Green), Honeybee (Yellow), Sheep (Orange), and Smelt (Purple).

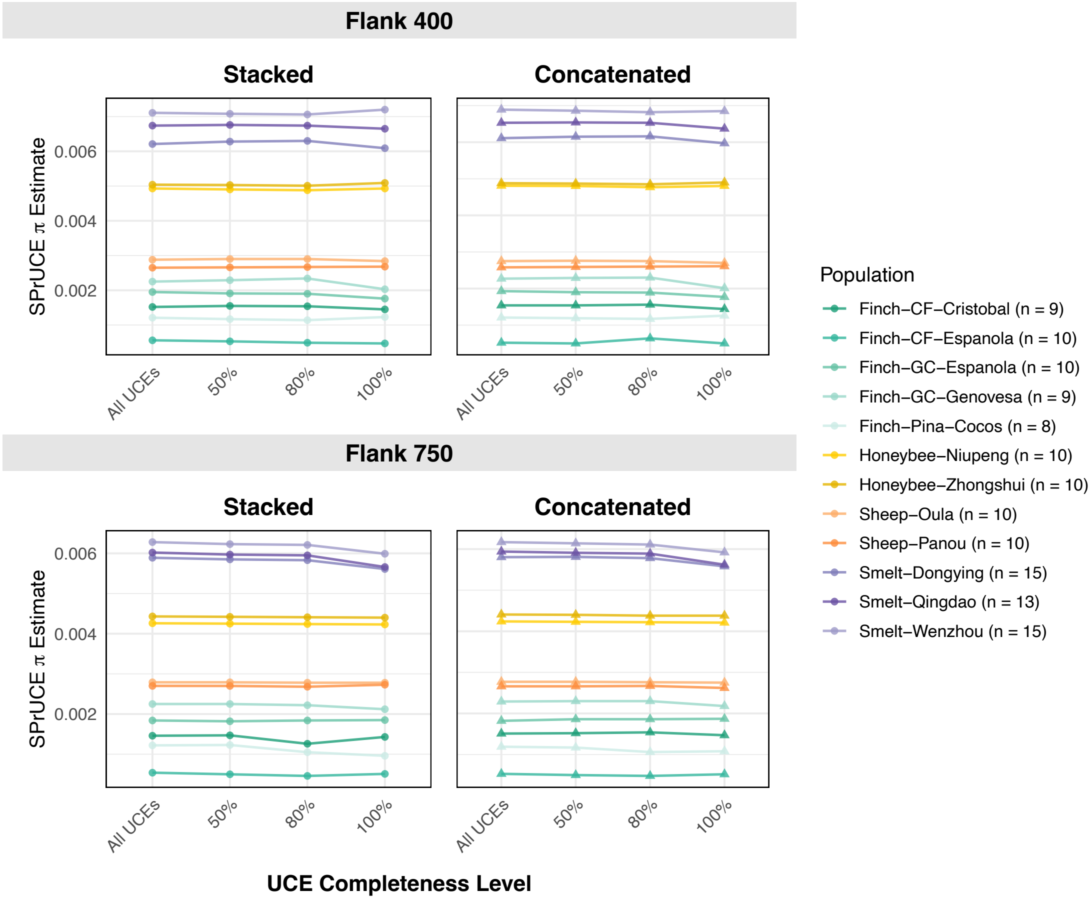

Figure S6: Estimated nucleotide diversity ($\pi$) across dataset completeness thresholds for all populations. Completeness thresholds were defined based on the minimum proportion of individuals required per UCE locus: All UCEs (no filtering), 50% (at least half of individuals present per locus), 80%, and 100% (all individuals present in every locus). The top panels show results for 400 bp flanking regions, and the bottom panels show results for 750 bp flanking regions. Each flank length includes estimates from the SPrUCE Stacked mode (left) and Concatenated mode (right). Population sample sizes ($n$) are shown in the legend.

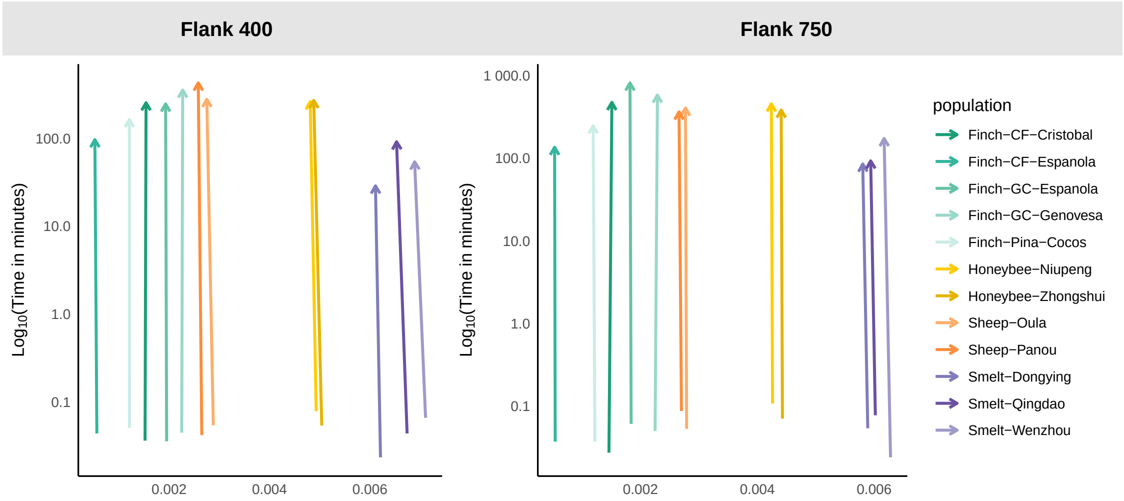

Figure S7: Comparison of runtime ($y$-axis) and nucleotide diversity estimates ($x$-axis) across methods and populations. Showing the relationship between log-transformed run time (in minutes) for Stacked to Concatenated methods. Results are grouped by flank length group: 400 bp flank and 750 bp flank.

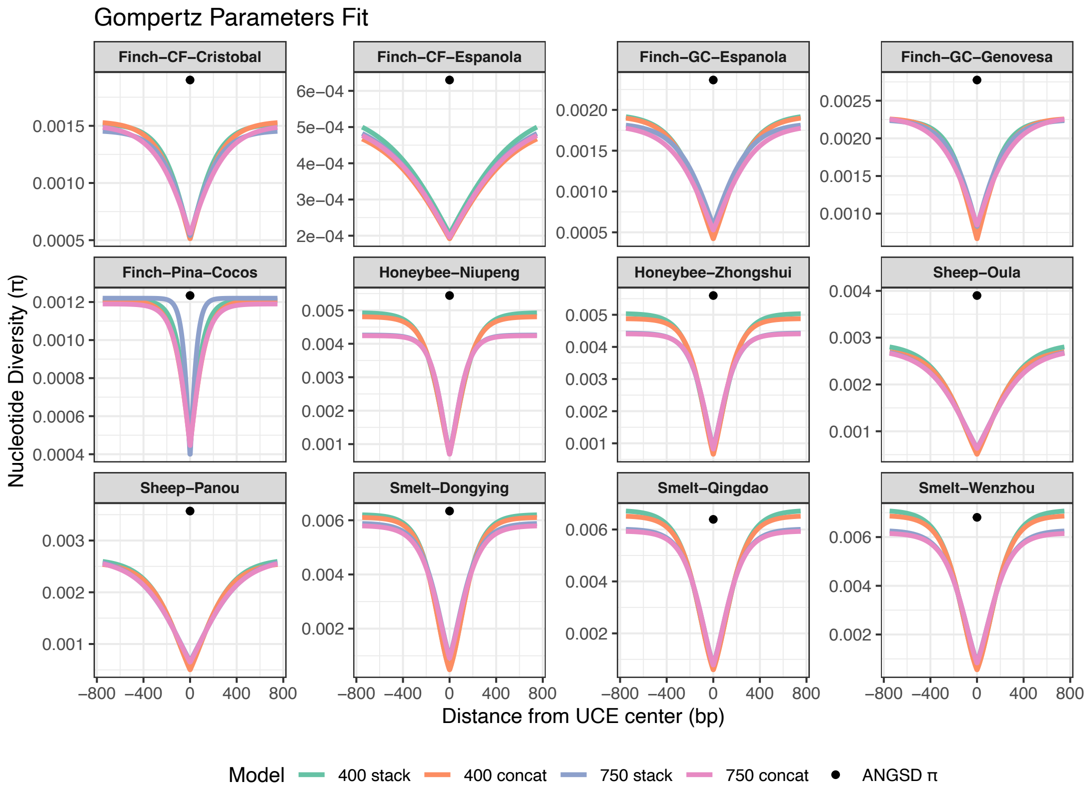

| Species Group | Population | Size | Stacked 400 | | Concat 400 | | Stacked 750 | | Concat 750 | |
|---|---|---|---|---|---|---|---|---|---|---|
| | | $n$ | $\beta$ | $\gamma$ | $\beta$ | $\gamma$ | $\beta$ | $\gamma$ | $\beta$ | $\gamma$ |
| *A. cerana* | Niupeng | 10 | 1.855 | 0.00959 | 1.915 | 0.01032 | 1.806 | 0.01284 | 1.839 | 0.01307 |
| | Zhongshui | 10 | 1.943 | 0.00936 | 2.030 | 0.01037 | 1.661 | 0.01031 | 1.704 | 0.01067 |
| *C. fusca* | Cristobal | 9 | 1.089 | 0.00733 | 1.110 | 0.00658 | 1.004 | 0.00696 | 1.000 | 0.00552 |
| | Espanola | 10 | 1.007 | 0.00291 | 1.002 | 0.00298 | 1.011 | 0.00292 | 1.001 | 0.00299 |
| *G. conirostris* | Espanola | 10 | 1.492 | 0.00592 | 1.534 | 0.00604 | 1.149 | 0.00577 | 1.245 | 0.00520 |
| | Genovesa | 9 | 1.200 | 0.00793 | 1.237 | 0.00717 | 1.005 | 0.00675 | 1.000 | 0.00553 |
| *P. inornata* | Cocos island | 8 | 1.021 | 0.01449 | 1.000 | 0.01190 | 1.160 | 0.02647 | 1.000 | 0.01261 |
| *O. aries* | Oula | 10 | 1.700 | 0.00549 | 1.700 | 0.00593 | 1.478 | 0.00480 | 1.486 | 0.00487 |
| | Panou | 10 | 1.645 | 0.00583 | 1.650 | 0.00612 | 1.402 | 0.00447 | 1.420 | 0.00468 |
| *S. sinica* | Dongying | 15 | 2.543 | 0.00980 | 2.572 | 0.00995 | 1.852 | 0.00895 | 1.875 | 0.00892 |
| | Qingdao | 13 | 2.364 | 0.00837 | 2.418 | 0.00888 | 2.030 | 0.00885 | 2.054 | 0.00908 |
| | Wenzhou | 15 | 2.513 | 0.00817 | 2.554 | 0.00860 | 1.996 | 0.00803 | 2.029 | 0.00833 |

Figure S8: The fitted Gompertz function across species and populations. Estimates of $\beta$ and $\gamma$ reflect the shape and decay rates of diversity decline with distance from UCE centers for both 400 bp and 750 bp flanking regions. The parameters estimated in each case is shown in the table below. Note how low values of $\gamma$ correspond to slower rates of reaching the asymptote, and coincide with higher deviations from the ANGSD estimates.

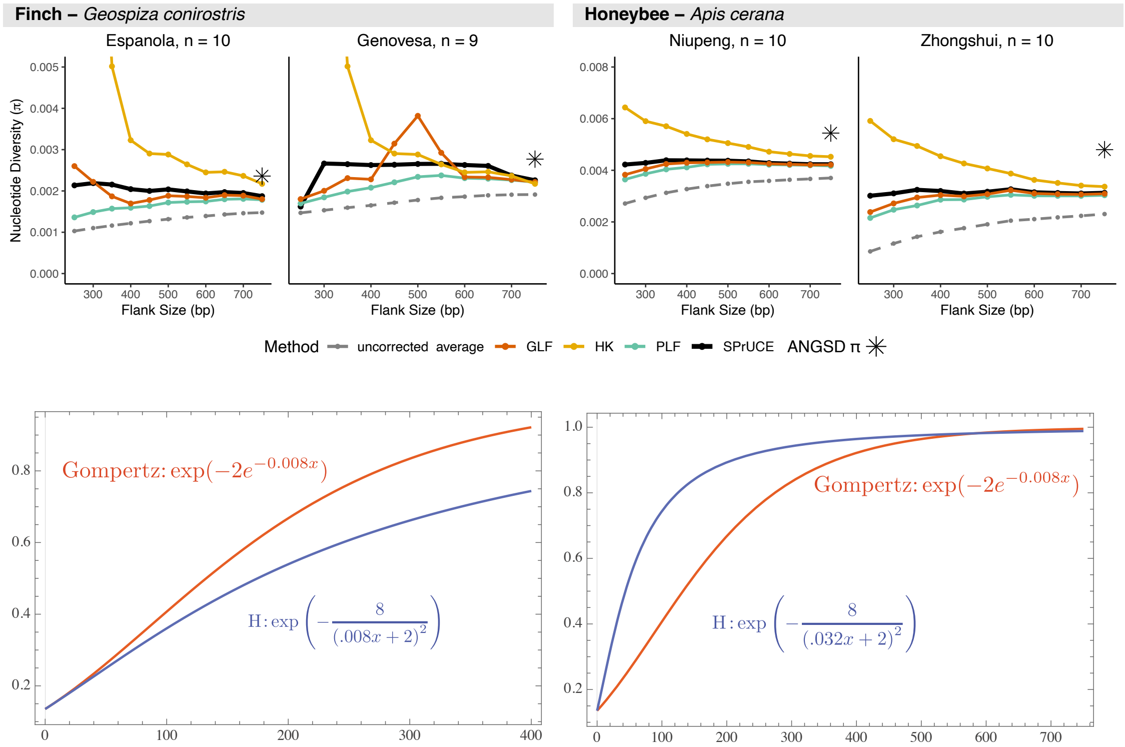

Figure S9: Top: Using the Hudson Kaplan equation ($\exp\left(-\frac{u\cdot s}{(2r\cdot x+2s)^2}\right)$) for modeling change in diversity versus other functions, including Gompertz. HK is far more sensitive to flank size; reducing the flank below 600bp can lead to very high estimates. Lack of robustness of HK discouraged us to use it, despite its theoretical appeal. Bottom: The HK equation has a slower rate of reaching its asymptote compared to Gompertz with equivalent parameters that, according to calculations, make the two functions approximate each other (left). The approximation is only good for small $x$. Given larger flank (right), if we chose parameters that force HK to get near the asymptote at about 750bp (by increasing the recombination rate $r$ four-fold), the HK model will predict a faster increase in diversity close to the core compared to Gompertz. Note that HK equation is convex, while Gompertz has an inflection point.

### S3 Supplementary text

#### S3.1 SPrUCE Algorithm

**procedure** ESTIMATEDIVERSITY
  **for** each UCE alignment $A_j$ in alignments **do**
    $i \leftarrow 0$
    **for** each position in the alignment $A_j$ **do**
      Identify major allele as the most frequent base (consensus)
      Categorize deviations as substitutions, insertions, or deletions
      **if** $|\text{non-gap alleles}| \leq 2$ **then**
        $i \leftarrow i + 1$
        $n_{i,j} \leftarrow$ Number of non-gap sequences in the column
        $s_{i,j} \leftarrow$ Number of times the major allele appears in the column
        $\pi_{i,j} \leftarrow$nucleotide diversity at position $i$ computed using $\frac{s_{i,j} \cdot (n_{i,j} - s_{i,j})}{\binom{n_{i,j}}{2}}$
      **end if**
    **end for**
  **end for**
  **if** mode = `concat` **then**
    Estimate $(\theta, \beta, \gamma)$ by minimizing

$$\sum_{j=1}^{k} \sum_{i=-\ell}^{\ell} \frac{n_{i,j} - 1}{n_{i,j} + 1} \left( f(i; \theta, \beta, \gamma) - \pi_{i,j} \right)^2 \ .$$

  **else** ▷ mode = `stack`
    $\forall i : n_i = \sum_j n_{i,j}$
    $f \leftarrow$ A spline function fit to the log of empirical cumulative distribution of $\log(n_i)$.
    $\ell' \leftarrow$ The inflection point of $f$ where its second derivative changes sign.
    Estimate $(\theta, \beta, \gamma)$ by minimizing

$$\sum_{i=-\ell'}^{\ell'} \frac{n_i - 1}{n_i + 1} \left( f(i; \theta, \beta, \gamma) - \frac{1}{k} \sum_{j=1}^{k} \pi_{i,j} \right)^2$$

  **end if**
  **return** The diversity estimates $(\theta)$ and other Gompertz parameters $(\beta, \gamma)$
**end procedure**

Algorithm S1: **SPrUCE.** Algorithm

#### S3.2 Approximation of $\pi_i$ by $2f_i$

1. For a site $i$, let $s_i$ be the count of individuals with the derived allele at site $i$, and $n$ be the number of individuals.

2. Let $f_i$ be the fraction of alleles that are one at position $i$.

3. Tajima's $\pi$ is estimated using by averaging the following quantity across sites:

$$\pi_i = \frac{s_i \cdot (n - s_i)}{\binom{n}{2}} \tag{S1}$$

$$\simeq 2\frac{s_i(n - s_i)}{n \cdot n} \tag{S2}$$

$$= 2f_i(1 - f_i) \tag{S3}$$

$$= 2f_i - 2f_i^2 \tag{S4}$$

$$\approx 2f_i \tag{S5}$$

   The approximation in Equation (S2) follow from $n^2 \gg n$ for a substantially large $n$ and the approximation in Equation (S5) follows from $f_i \gg f_i^2$ for a small value of $f_i$, such as 0.01 or lower.

4. The `phyluce_align_get_smilogram_from_alignments` function in Phyluce outputs $f_i$ for each position along the UCE, stacking all positions $i$ of all UCEs on top of one another, effectively making $n$ to be the number of individuals multiplied by the number of loci, which is often a very large number.

5. Calculations above show that if the "substitution frequency" quantity outputted by Phyluce is to be interpreted as genetic diversity, and if we decide to ignore the impact of selection, we still need to multiply the "substitution frequency" by a factor of 2.
